# P2Y12–P-Selectin Mediated Platelet Activation Drives Dengue-Associated Thrombocytopenia

**DOI:** 10.1101/2025.09.08.674864

**Authors:** Viviane Lima Batista, Jenniffer Ramos Martins, Angélica Samer Lallo dias, Talita Cristina Martins da Fonseca, Letícia Soldati Silva, Felipe Emanuel Oliveira Rocha, Pedro Augusto Carvalho Costa, Wallison Nunes da Silva, Jessica Aparecida Barsalini Pereira, Maisa Antunes, Pedro Pires Goulart Guimarães, Gustavo Batista de Menezes, Adriano de Paula Sabino, Lirlândia Pires Sousa, Celso Martins Queiroz-Junior, Eugenio Damaceno Hottz, Mauro Martins Teixeira, Vivian Vasconcelos Costa

**Affiliations:** Department of Microbiology; Department of Biochemistry and Immunology; Department of Morphology; Department of Pharmacology, Institute of Biological Sciences, Universidade Federal de Minas Gerais, Belo Horizonte, Brazil; Laboratory of Clinical, Experimental and Molecular Hematology, Department of Clinical and Toxicological Analysis Faculty of Pharmacy; Department of Clinical and Toxicological Analysis, Universidade Federal de Minas Gerais, Belo Horizonte, Brazil; Laboratory of Immunothrombosis, Department of Biochemistry, Universidade Federal de Juiz de Fora (UFJF), Juiz de Fora, Brazil

## Abstract

Dengue virus (DENV) infection frequently causes thrombocytopenia, a hallmark of severe disease. The underlying mechanisms of this condition remain incompletely understood. Using a murine model that mirrors human hematologic and inflammatory responses, we show that DENV impairs megakaryopoiesis, causing necrotic loss of bone marrow megakaryocytes and compensatory thrombopoietin elevation. Megakaryocyte-derived platelets exhibit mitochondrial dysfunction and phosphatidylserine exposure, consistent with apoptosis. Systemically, DENV drives P-selectin– dependent platelet activation and platelet–monocyte aggregates, accompanied by increased plasmatic chemokines (CXCL4, CCL5, CXCL1) and vascular leakage in target organs such as liver and lungs, reflecting heightened thromboinflammation. Pharmacologic blockade of P-selectin or inhibition of P2Y12 with clopidogrel restored platelet counts, rescued megakaryocyte numbers, and reduced systemic inflammation. Collectively, these findings demonstrate for the first time in an *in vivo* model that DENV-induced thrombocytopenia arises from combined megakaryocyte impairment and platelet hyperactivation and highlight platelet-targeted interventions as potential therapeutic strategies.

**KEY POINTS:**

- DENV disrupts platelet production and survival via megakaryocyte necrosis and intramedullary platelet apoptosis, driving thrombocytopenia.
- DENV triggers thrombocytopenia via P-selectin platelet activation; blocking it with antibody P-selectin or clopidogrel reduces inflammation.

## 1. INTRODUCTION

Dengue is a mosquito-borne viral disease caused by four antigenically distinct dengue virus serotypes (DENV-1 to -4) and primarily transmitted by *Aedes aegypti* [1]. Its rapid global spread and high incidence make dengue a major public health concern [1–3]. Clinical manifestations range from mild disease without warning signs to severe dengue, which can result in life-threatening complications such as thrombocytopenia, vascular leakage, hemorrhage, multi-organ failure, hypovolemic shock, and death [4,5]. Endothelial dysfunction and thromboinflammation are central to dengue pathogenesis and represent potential targets for therapeutic intervention.

Thrombocytopenia is a defining feature of dengue, with platelet counts frequently dropping below 50,000/μL in severe cases and coinciding with hemorrhagic manifestations [6–8]. Although the mechanisms driving DENV-induced thrombocytopenia remain incompletely defined, current evidence points to impaired platelet production in the bone marrow together with increased peripheral destruction and clearance. [9]. Several studies have demonstrated a positive correlation between platelet activation and thrombocytopenia during DENV and other infections, such as SASRCOV-2 [4,10–12].

Platelets from dengue patients display heightened activation, evidenced by increased surface expression of P-selectin and binding of PAC-1 antibody to the activated αIIbβ3 integrin complex. [4,13,14]. The formation of platelet–leukocyte aggregates is a key driver of thromboinflammation and vascular injury in severe dengue [15]. This process is mediated by P-selectin translocation to the platelet surface, facilitating interactions with leukocytes and endothelial cells via PSGL-1 (P-selectin glycoprotein ligand-1) [16,17]. Activated platelets amplify the inflammatory response by releasing pro-inflammatory and procoagulant mediators, such as platelet factor 4 (PF-4/CXCL4), neutrophil-activating peptide 2 (NAP-2), and platelet-activating factor (PAF), thereby contributing to enhanced inflammation and thrombosis[18]. Endothelial P-selectin expression further enhances interactions with PSGL-1–expressing leukocytes, amplifying this pathogenic cascade [16,17]. We hypothesize that P-selectin is a central mediator of thromboinflammation during dengue virus (DENV) infection. Supporting this, murine models of sickle cell disease have shown that blocking P-selectin with anti-CD62P monoclonal antibodies reduces platelet–neutrophil aggregates and mitigates lung injury. Indeed, P-selectin-deficient mice (P[/[) display reduced platelet–leukocyte aggregation and thrombus formation in models of deep vein thrombosis, highlighting the essential role of P-selectin in these processes [19–21]. In this study, we investigated the mechanisms underlying dengue virus-induced thrombocytopenia using an *in vivo* model in interferon-alpha/beta receptor knockout mice (A129 strain). These mice exhibit high susceptibility to DENV infection and develop macroscopic and histopathological alterations that closely resemble key features of the disease in humans. [22,23] . We demonstrate that dengue infection causes progressive thrombocytopenia through bone marrow megakaryocyte death, platelet–leukocyte aggregates, and elevated P-selectin in blood and target organs. P-selectin blockade reversed these effects, establishing its central role in dengue-associated thromboinflammation and pathogenesis.

## 2. METHODS

### 2.1 Mice

7-9-week-old male and female A129 mice, equally distributed, originally obtained from the Bioterium at the University of São Paulo (USP) and bred at the Bioterium of the Department of Biochemistry and Immunology, Federal University of Minas Gerais (ICB/UFMG) were used. All procedures complied with institutional guidelines and were approved by the UFMG Animal Ethics Committee (CEUA/UFMG, protocol 12/2023). Mice were housed in microisolator cages under BSL-2 conditions, with controlled temperature (24 ± 2°C), a 12-hour light/dark cycle, and *ad libitum* access to food and water.

### 2.2 Cell Lines, Monoclonal Antibodies, and Virus

Vero CCL81 (BCRJ 0245) and *Aedes albopictus* C6/36 (BCRJ 0343) cells were obtained from the Rio de Janeiro Cell Bank (BCRJ) and grown in DMEM or L15 medium (Cultilab), respectively, both supplemented with 10% heat-inactivated fetal bovine serum. For *in vivo* studies, a low-passage DENV-2 clinical isolate (strain 3295, GenBank: EU081177.1) was amplified in C6/36 cells, and the supernatants were collected, filtered, concentrated, titrated by plaque assay, and stored at –80°C until use.

### 2.3 Infection and Treatment

Mice were infected subcutaneously in the right hind paw (intraplantar route) with 30 μL of RPMI medium containing (2×10[, 2×10^3^ or 2×10^2^ PFU of DENV-2). Control (Mock) mice received the same volume of cell culture supernatant. Mice were monitored daily for clinical signs and mortality, defined as death or weight loss ≥ 20% of baseline, and euthanasia was performed on days 1, 3, and 5 post-infections. For P-selectin analysis, one hour before infection, mice received an intraperitoneal injection containing 30 µg of anti–P-selectin (CD62P) antibody (R&D Systems, MAB737) in 200 μL of saline as described previously[19] . Control groups received either saline alone or 30 µg of human IgG isotype control (Sigma, 1001709695). The antibody was administered again at 48 hours post-infection at the same dose, and mice euthanized 3 days after the infection. For Clopidogrel treatment, we followed the protocol from An et al. (2018)[24]. One hour after the infection, Clopidogrel bisulfate (Sigma, SML0004) was given by gavage in a 0.5% carboxymethylcellulose solution (Synth). A loading dose of 15 mg/kg was given on day 1, followed by a daily maintenance dose of 5 mg/kg for the next three days. Control mice received vehicle solutions.

### 2.4 Sample Collection

At specific time-points, mice were anesthetized with ketamine (90 mg/kg) and xylazine (10 mg/kg), and blood was collected from the vena cava using EDTA-coated syringes. Platelets were counted by diluting the blood 1:1000 in ammonium oxalate and analyzing it in a Neubauer chamber. Results were expressed as platelet count × 10³ per mL of blood. The left liver and lung lobes, right femur, and right paw were fixed in 4% buffered formaldehyde for histology, immunohistochemistry, and mast cell degranulation analysis. The right lung lobes, brain, liver, and spleen were frozen in liquid nitrogen and stored at -80°C for viral load, ELISA, and other tests. In additional experiments, mice were used to evaluate vascular permeability as described previously [25]. In another group, whole blood, femur, and lungs were collected for flow cytometry analysis.

### 2.5 Flow Cytometry Assays

Immunophenotyping of infiltrating cells, platelet profile, and platelet–leukocyte aggregates was performed using whole blood, bone marrow, and lung tissue. Red blood cells were lysed with ACK Lysing Buffer. Lung tissue was digested with collagenase type III (3 mg/mL) for 40 minutes at 37°C, dissociated, filtered, and lysed to remove red blood cells. Antibodies are listed in Table S1. Cells were stained with three panels: (i) lymphoid cells with platelet aggregates, (ii) myeloid cells with aggregates and megakaryocytes, and (iii) platelet production and activation. A separate experiment assessed megakaryocyte and platelet death. Absolute cell quantification was performed using Count Bright™ spheres (ThermoFisher). Dead cells were excluded with Live/Dead reagent (Invitrogen). Flow cytometry analysis included debris exclusion, doublet removal (FSC-A vs. FSC-H), and elimination of abnormal time vs. FSC-A events to minimize artifacts. Only live CD45+ cells were analyzed, and subpopulations were identified by specific markers **(Table S1)**. Platelet profile was measured on a logarithmic scale to exclude debris, leukocytes, and megakaryocytes. CD41+ expression was quantified in subsets from pulmonary and peripheral compartments. Data were analyzed with FlowJo V10.4.11 (BD) and GraphPad Prism V7.0 (GraphPad). The complete gating strategy is provided in **Supplementary Figure 2**.

### 2.6 Vascular Permeability and Confocal Intravital Microscopy

Evans blue dye extravasation into the lungs and spleen was used as an indicator of increased vascular permeability, as described previously [25,26]. The amount of Evans blue in the tissue was determined by comparing the absorbance measured against a standard curve of the dye, read at 620 nm in a spectrophotometric plate reader. Results are expressed as the amount of Evans blue per 100 mg of tissue. Vascular permeability in the liver was assessed by intravenously injecting 100 μL of albumin-FITC (5 mg/mL, Sigma-Aldrich®) into the lateral tail vein of mice. After 30 minutes, the animals were anesthetized intraperitoneally (i.p) with a mixture of ketamine (90 mg/kg) and xylazine (10 mg/kg) for laparotomy, following the protocol previously described Marques et al. (2015) [27]. After surgery, the animals were positioned on an acrylic support adapted for microscopy, with the left lateral liver lobe placed on the slide, allowing tissue imaging using the NIS-Elements Viewer software. Vascular extravasation was then analyzed by quantifying fluorescence in the FIJI image analysis program. Platelet and leukocyte aggregates in the lungs were assessed by intravenously injecting monoclonal antibodies anti-CD45 (HI30, APC, eBioscience™, 6 μg/mouse) and anti-CD41 (PE, eBioscience™, 6μg/mouse), following the same procedure described previously [27].

### 2.7 Viral Titers Determination

Viral titration was measured in the hind paw, plasma, spleen, bone marrow, liver, lung, and brain. Blood was collected in EDTA tubes and centrifuged at 3,000×g for 15 minutes at room temperature. The plasma was then stored at -80°C. Organs were collected at different time points, weighed, and frozen at -80°C. Later, they were homogenized in RPMI 1640 medium (without FBS) to make 10% tissue suspensions. Virus titers in both the tissue homogenates and plasma were measured using a plaque assay with Vero cells. Results were reported as plaque-forming units (PFU) per gram of tissue or per mL of plasma. The minimum detectable level was 100 PFU per gram of tissue or per mL of plasma. [23].

### 2.8 Histology, Immunohistochemistry, and *In Vivo* Mast Cell Degranulation

Lung and liver samples were fixed in formaldehyde, paraffin-embedded, sectioned (5 μm), and stained with H&E. Inflammatory scores were blindly evaluated by a pathologist (CMQJ) following established protocols [28,29]. Femur samples were decalcified with EDTA 10% for 30 days and subsequently processed for embedding in paraffin. Sections of the femur from infected and control groups were qualitatively analyzed in the bone marrow for replicating viral detection using immunohistochemistry for anti-dsRNA (mouse monoclonal antibody detecting double-stranded RNA; Millipore, 1:100) manufacturer’s instructions. For mast cell degranulation analysis, the right hind paw plantar pad was collected and processed histologically. Mast cells were stained with toluidine blue and classified as degranulated or normal based on their morphological characteristics, following a previously described protocol [30].

### 2.9 Inflammatory Molecules Quantification by Elisa

Plasma and tissue levels of CCL2, CXCL1, CCL5, and CD62P were measured by ELISA using the DuoSet kit (R&D Systems). Plasma MCPT-1 was assessed with the Ready-SET-Go kit (eBioscience). PF4/CXCL4 and thrombopoietin concentrations were determined using the Quantikine ELISA kit (R&D Systems).

### 2.10 Statistical Analyses

Statistical analyses were done using GraphPad Prism software (v.8.0.2). Data normality was tested with the Shapiro-Wilk test. For normally distributed data, a 95% confidence interval was used, and two-way ANOVA followed by Dunnett’s multiple comparisons test was applied. For non-normal data, a 99% confidence interval was used, and the Kruskal-Walli’s test followed by Dunn’s post-test was applied.

## 3. RESULTS

### 3.1 Subcutaneous Dengue Virus Infection in A129 Mice Recapitulates Progressive Thrombocytopenia and Disease

To mimic natural infection and monitor the development of thrombocytopenia, A129 mice were inoculated intraplantarly with 30 μL of dengue virus serotype 2 (DENV-2) **(Supplementary Figure 1A, Figure 1A)**. Infected mice displayed progressive weight loss from day 2 post-infection (dpi), along with clinical manifestations including hunched posture, conjunctivitis, ruffled fur, and reduced mobility (**Table S2, Supplementary Figure 1B; Figure 1B)**. All inoculated mice eventually succumbed to infection. The highest inoculum tested (2 × 10[PFU per mouse) caused death by day 8 and induced a severe and progressive disease phenotype. This inoculum was therefore selected for subsequent experiments **(Supplementary Figure 1C, Figure 1B)**. DENV-2 infection induced progressive thrombocytopenia **(Figure 1C)** and elevated systemic levels of the chemokines CCL2, CXCL1, and CCL5 in plasma and spleen **(Figure 1D)**. Viral replication was detected as early as 1-day post-infection (dpi) in the footpad, spleen, liver, and plasma, with subsequent dissemination to the brain at later time points **(Figure 1E).** Infection also promoted mast cell degranulation, evidenced by increased plasma concentrations of MCPT-1 and higher numbers of degranulated mast cells on days 3 and 5 dpi **(Figure 1F).** Inflammatory infiltrates were observed in the liver **(Figure 1G)**, accompanied by increased hepatic vascular permeability **(Figure 1H).** Splenomegaly was prominent **(Supplementary Figure 1D–E)** and was associated with increased splenic vascular permeability **(Supplementary Figure 1F)**. Together, these findings show that A129 mice are highly susceptible to subcutaneous DENV-2 infection and develop a severe, systemic, and lethal disease that recapitulates key features of severe human dengue.

**Figure 1:**
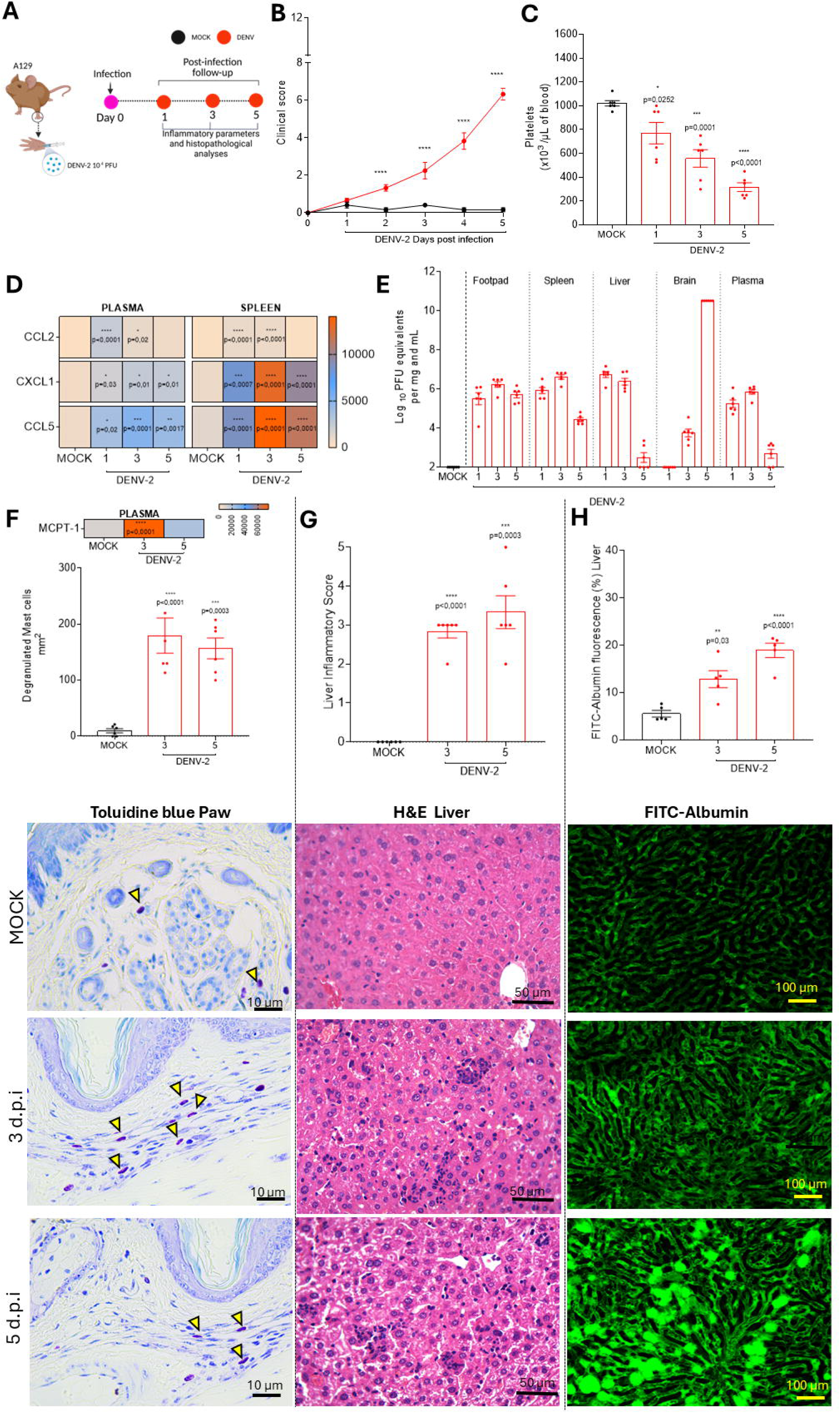
Characterization of disease parameters in A129 mice infected with a clinical isolate of DENV-2. (A) Experimental scheme: A129 mice (n = 6 per group), males and females, equally distributed, were inoculated subcutaneously with DENV-2 at a concentration of 2×10LJ PFU/animal or with cell culture supernatant (Mock group). Mice were monitored daily for weight loss and clinical score, and were euthanized on days 1, 3, and 5 post-infections for blood and tissue collection for subsequent analyses. **(B) Clinical score: Mice** were assessed daily for disease severity. Clinical score was determined based on the following parameters: weight loss, spinal hunching, diarrhea, ocular inflammation, decreased locomotor activity, and prostration. Score was assigned based on severity: 0 (absence of symptoms), 0.5 (mild), 1 (moderate), and 2 (severe). The graph shows the sum of the scores assigned, as explained in Table S2. **(C) Platelet count:** The manual quantification of platelets in peripheral blood was performed using a Neubauer chamber, with results expressed as 10³ platelets µl of blood. **(D) Inflammatory mediator heatmap:** The quantification of the inflammatory mediators CCL2, CXCL1, and CCL5 in plasma and spleen was performed by ELISA, with results expressed in pg/mL of plasma or pg/mg of splenic tissue. **(E) Viral titer:** Determined in organ extracts (paw, spleen, liver, and brain) and plasma of DENV-2 infected mice through plaque assay. Results are expressed as Log10 PFU per gram of tissue or per milliliter of plasma. **(F) Analysis of mast cell degranulation:** degranulated mast cell count and ELISA assay for MCPT-1. Manual counting of degranulated mast cells in the paw and representative images of histological sections of the plantar pad stained with toluidine blue, obtained on days 3 and 5 post-infection. **(G) Liver analyses:** Inflammatory infiltrate in the liver: Evaluation of the presence of hepatic inflammatory infiltrate. Representative images of histological sections of the liver stained with hematoxylin and eosin (H&E). **(H) Confocal microscopy of the liver:** Vascular permeability in the liver: Evaluation of hepatic vascular permeability by confocal microscopy, with data expressed as the percentage of FITC-albumin fluorescence (n=5 per group) Representative images obtained by confocal microscopy. Evaluation of hepatic vascular permeability by confocal microscopy, with data expressed as the percentage of FITC-albumin fluorescence (n=5 per group). Symbols (*, **, ***, ****) indicate statistically significant differences between the infected groups and the control (Mock) group (p < 0.05), as determined by one-way ANOVA followed by Dunnett’s post-test. Figure A was created with Biorender. Batista, VL (2025).

### 3.2 DENV-infection triggers megakaryocyte’s death and platelet impairment in bone marrow

Decreased megakaryocyte and platelet numbers, along with impaired platelet function, are frequently observed in dengue patients, and direct megakaryocyte infection has been proposed as a contributing mechanism to dengue-associated thrombocytopenia [8]. In our model, dengue virus infected bone marrow megakaryocytes, as evidenced by increased viral titers **(Figures 2B)** and positive staining for double-stranded RNA (dsRNA), indicative of active replication **(Figure 2C).** Infection was accompanied by a significant reduction in megakaryocyte numbers **(Figure 2D; Supplementary Figure 2A)**. Flow cytometry revealed that ∼7% of megakaryocytes exhibited apoptotic markers (CD45[CD41[Annexin V[PI[) **(Figure 2E)**, while ∼40% displayed necrotic markers (CD45[CD41[Annexin V[PI[) **(Figures 2F).** Bone marrow flushes showed a notable increase in platelet percentages post-infection **(Supplementary Figure 2B; Figure 2G)**, and by day 3, newly released platelets demonstrated activation, with elevated surface P-selectin expression **(Figure 2H)**. On days 1 and 5, these platelets exhibited mitochondrial membrane depolarization (TMRE loss) and increased Annexin V binding, consistent with apoptosis **(Figures 2I–J)**. These findings show that DENV infects bone marrow megakaryocytes, leading to their impairment and the release of platelets with signs of activation and apoptosis into the marrow environment.

**Figure 2:**
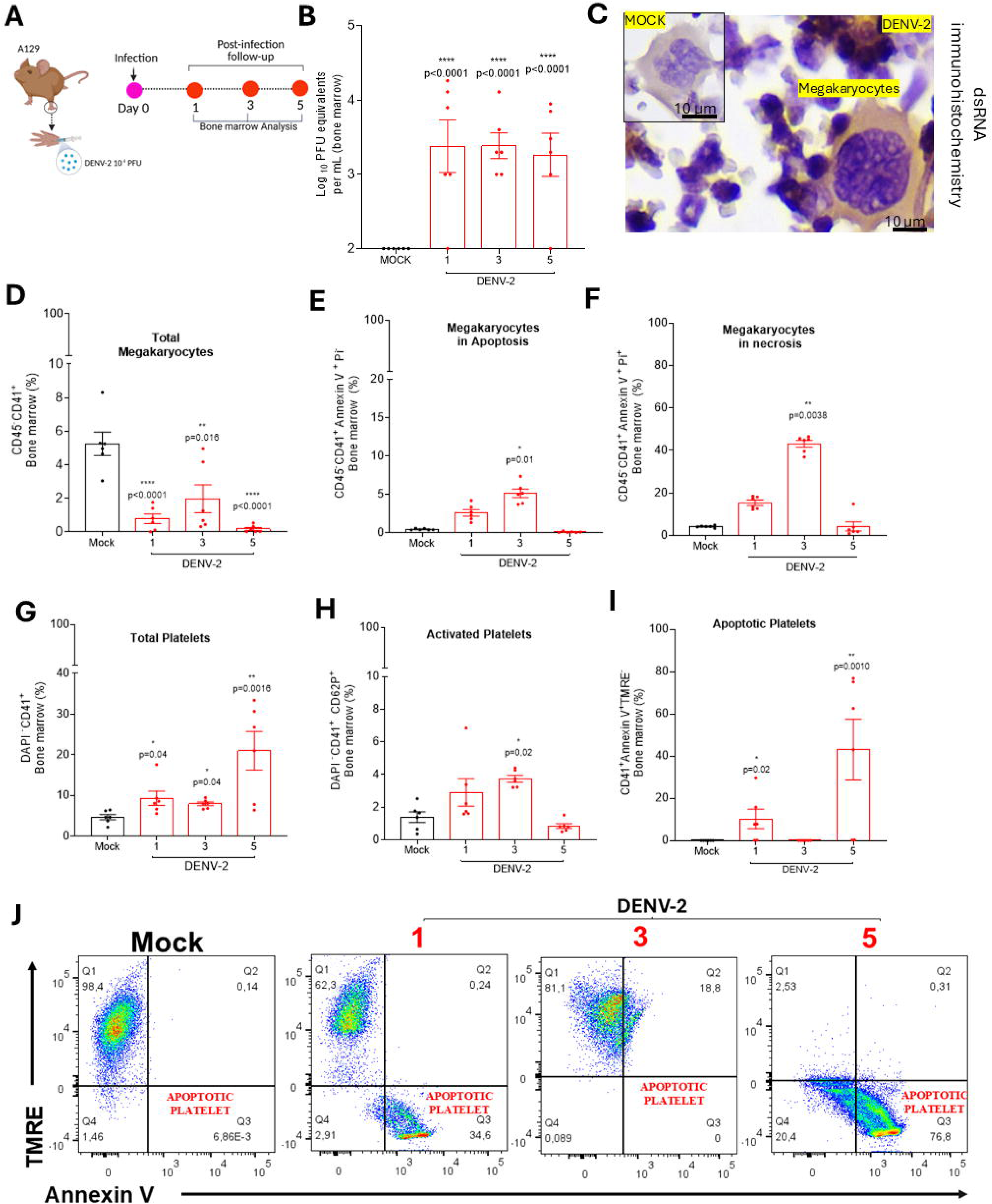
Characterization of Megakaryocyte and Platelet Profiles in Bone Marrow After Dengue Virus Infection. (A) Experimental scheme: A129 mice (n = 6 per group) were inoculated subcutaneously with DENV-2 (2×10LJ PFU/animal) or with cell culture supernatant (Mock group). Mice were monitored daily for weight loss and clinical score and were euthanized on days 1, 3, and 5 post-infections. After euthanasia, bone marrow from the femur of the right hind limb was collected for flow cytometry analysis, and from the left hind limb for histology and immunohistochemical analyses. **(B) Viral titration in bone marrow flush.** Viral titers were determined by plaque assay and are expressed as LogLJLJ PFU per gram of tissue or per milliliter of plasma. **(C) Immunohistochemistry for dsRNA:** Detection of dengue virus double-stranded RNA (dsRNA) using a specific probe in bone marrow from uninfected control (Mock) and DENV-infected mice. **(D) Percentage of megakaryocyte**: quantification in bone marrow: Positive selection of CD45^-^CD41 cells by flow cytometry. **(E)Percentage of megakaryocytes in apoptosis:** The CD45^-^CD41LJLJ ANEXIN VLJPI^-^ population was considered. **(F) Percentage of megakaryocytes in necrosis:** The CD45^-^CD41LJANEXIN VLJPILJ population was considered**. (G) Percentage of platelets.** Considering the DAPILJCD41LJ population. **(H) Percentage of Activated Platelet. (I) Percentage of apoptotic platelets:** The CD41LJANEXIN VLJ population was considered, with loss of mitochondrial membrane potential assessed by TMRM dye. **(J) Representative dot plot of the platelet gating strategy with apoptosis staining**. All platelet data were analyzed on a logarithmic scale to exclude larger cells such as megakaryocytes. Symbols (*, **, ***, ****) indicate statistically significant differences between infected and control (Mock) groups (p < 0.05), as determined by one-way ANOVA followed by Dunnett’s post-test. Figure A was created with Biorender. Batista, VL (2025).

### 3.3 Peripheral Platelet Activation Promotes Platelet-Leukocyte Aggregation and Aggravates DENV-Induced Thrombocytopenia

Spearman analysis revealed a significant inverse correlation between bone marrow platelets exhibiting apoptotic markers and peripheral blood platelet counts (r = –0.4, p = 0.02) **(Supplementary Figure 1G)**, suggesting that premature platelet death may impair their release into circulation **(Figures 3A–B**). Significant platelet activation was detected on the third day post-infection, as indicated by increased surface expression of P-selectin **(Figures 3C–D)**. In parallel, soluble P-selectin levels were elevated as early as the first day of infection **(Figure 3E).** Infection also led to increased levels of the platelet-derived chemokine PF4/CXCL4 and elevated plasma thrombopoietin concentrations, observed on the 5th dpi. Furthermore, elevated levels of the viral NS1 protein were detected in the plasma at 3 dpi **(Figure 3E).** Platelet activation was accompanied by the formation of aggregates with neutrophils, monocytes, and dendritic cells **(Figures 3F–H, Supplementary Figure 2C-D**). At 1 dpi, platelet aggregates with CD4[T cells were observed **(Figures 3 K)**, whereas by 5 dpi, aggregates containing CD8[T cells and NKT cells became predominant **(Figures 3L, M)**. Taken together, these findings demonstrate that DENV-induced thrombocytopenia is driven by impaired thrombopoiesis in the bone marrow, along with increased platelet activation and interaction with leukocytes in peripheral blood.

**Figure 3:**
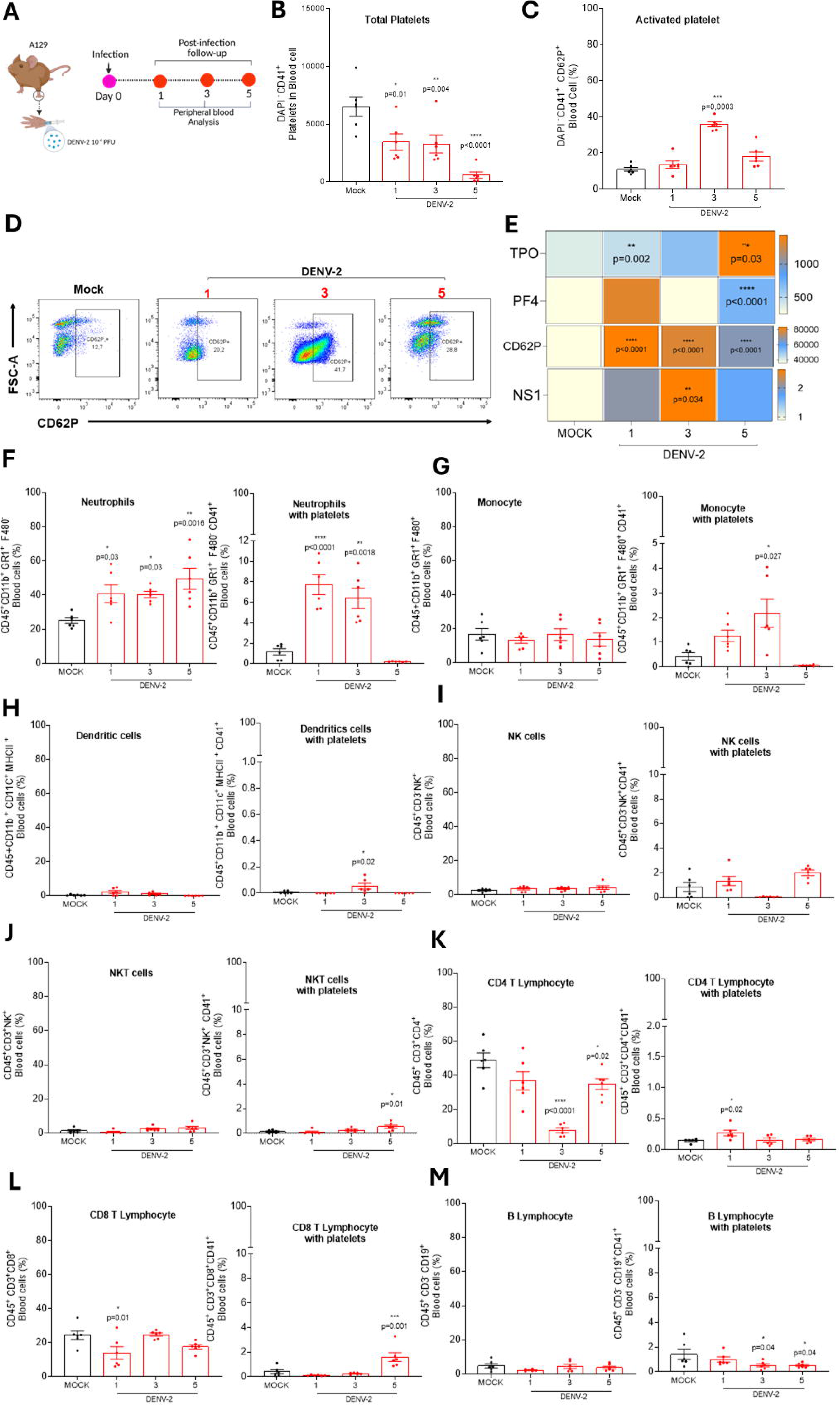
Evaluation of Dengue Virus-Induced Thrombocytopenia in Peripheral Blood. (A) Experimental design: A129 mice (n = 6 per group), were subcutaneously inoculated in the plantar region with DENV-2 at a concentration of 2×10LJ PFU/animal or with saline solution (Mock group). Mice were monitored daily for weight loss and clinical score and euthanized on days 1, 3, and 5 post-infections, followed by blood collection for flow cytometry analysis. **(B) Total platelets in peripheral blood:** Selected from the DAPILJ CD41LJ cell population**. (C) Activated platelets:** Selected based on the DAPILJ CD41LJ CD62PLJ population**. (D) Representative dot plots of platelet activation in peripheral blood. (E) Heatmap of plasma.** Thrombopoietin (TPO), PF4 (CXCL4), CD62P soluble and NS1 concentrations in plasma were measured by ELISA and expressed as pg/mL of plasma. **(F) Percentage of neutrophils:** Identified in the CD45LJ CD11bLJ GR-1LJ F4/80LJ population, including aggregates with CD41LJ platelets. **(G) Percentage of monocyte:** Selected from the CD45LJ CD11bLJ GR-1LJ F4/80LJ population and aggregated with CD41LJ platelets. **(H) Percentage of Dendritic cells:** Identified in the CD45LJ CD11bLJ CD11cLJ MHCIILJ population, including aggregates with CD41LJ platelets. **(I) Percentage of NK cells:** Selected from the CD45LJ CD3LJ NKLJ population and aggregated with CD41LJ platelets. **(J) Percentage of NKT cells:** Identified in the CD45LJ CD3LJ NKLJ population and aggregated with CD41LJ platelets. **(K) Percentage of CD4**D **T lymphocytes:** Selected from the CD45LJ CD3LJ CD4LJ population and aggregated with CD41LJ platelets. **(L) Percentage of CD8**D **T lymphocytes:** Selected from the CD45LJ CD3LJ CD8LJ population and aggregated with CD41LJ platelets. **(M) Percentage of B lymphocytes:** Selected from the CD45LJ CD3LJ CD19LJ population and aggregated with CD41LJ platelets. Data presented in Figures D-E were obtained through logarithmic scale reading in the cytometer, excluding larger leukocytes. Figure E was generated by multiplying the total frequency by the total platelet count in peripheral blood. The symbols (*, **, ***, ****) indicate statistically significant differences between infected groups and the control group (Mock) (p < 0.05). Statistical analysis: Figures B and G: Kruskal-Walli’s test followed by Dunn’s post-test. Figures C-F and H-L: One-way ANOVA followed by Dunnett’s post-test. Figure A was created with Biorender. Batista, VL (2025).

### 3.4 Lungs as a Refractory Target of DENV, Impairing Thrombopoiesis and Contributing to Platelet-Leukocyte Aggregation

Emerging evidence suggests that the lungs represent a significant site of platelet production, contributing 7–50% of total platelet output depending on physiological or experimental condition [31,32]. We next explored how dengue virus infection affects the lungs and whether this is associated with DENV-induced thrombocytopenia and megakaryocyte dynamics. Viral replication in the lungs peaked on day 3 after-infection, as evidenced by increased viral titers in the tissue **(Figure 4A-B).** Notably, a reduction in total megakaryocyte counts was observed on day 5 **(Figure 4C).** We then investigated the presence of platelet-leukocyte aggregates in the lungs. On day 3, flow cytometry of lung homogenates revealed that approximately 10% of lung cells were double-positive for CD45 and CD41, compared with the uninfected (mock) group **(Figures 4D).** This was further confirmed by confocal microscopy, which revealed the accumulation of CD45[CD41[cells in the pulmonary vasculature **(Figure 4E).** Platelets were found aggregated with neutrophil, macrophages, and dendritic cells **(Figures 4F-G, supplementary Figure 3A),** as well as with CD4 T lymphocyte cells on day 1 post-infection **(Supplementary Figures 3B)**. These platelet-leukocyte aggregates coincided with increased levels of CCL2, CXCL1, and CCL5 in lung tissue on day 3 **(Figure 4H)**, indicating a heightened inflammatory environment. This inflammation was associated with lung tissue damage **(Figures 4I-J)** and increased vascular permeability **(Figures 4K-L)**, indicating disruption of the pulmonary vascular barrier. These findings underscore the lungs as an important site of platelet production that is targeted during dengue virus infection. Infection disrupts megakaryocyte homeostasis and drives platelet-leukocyte aggregation, contributing to local inflammation and systemic thrombocytopenia.

**Figure 4:**
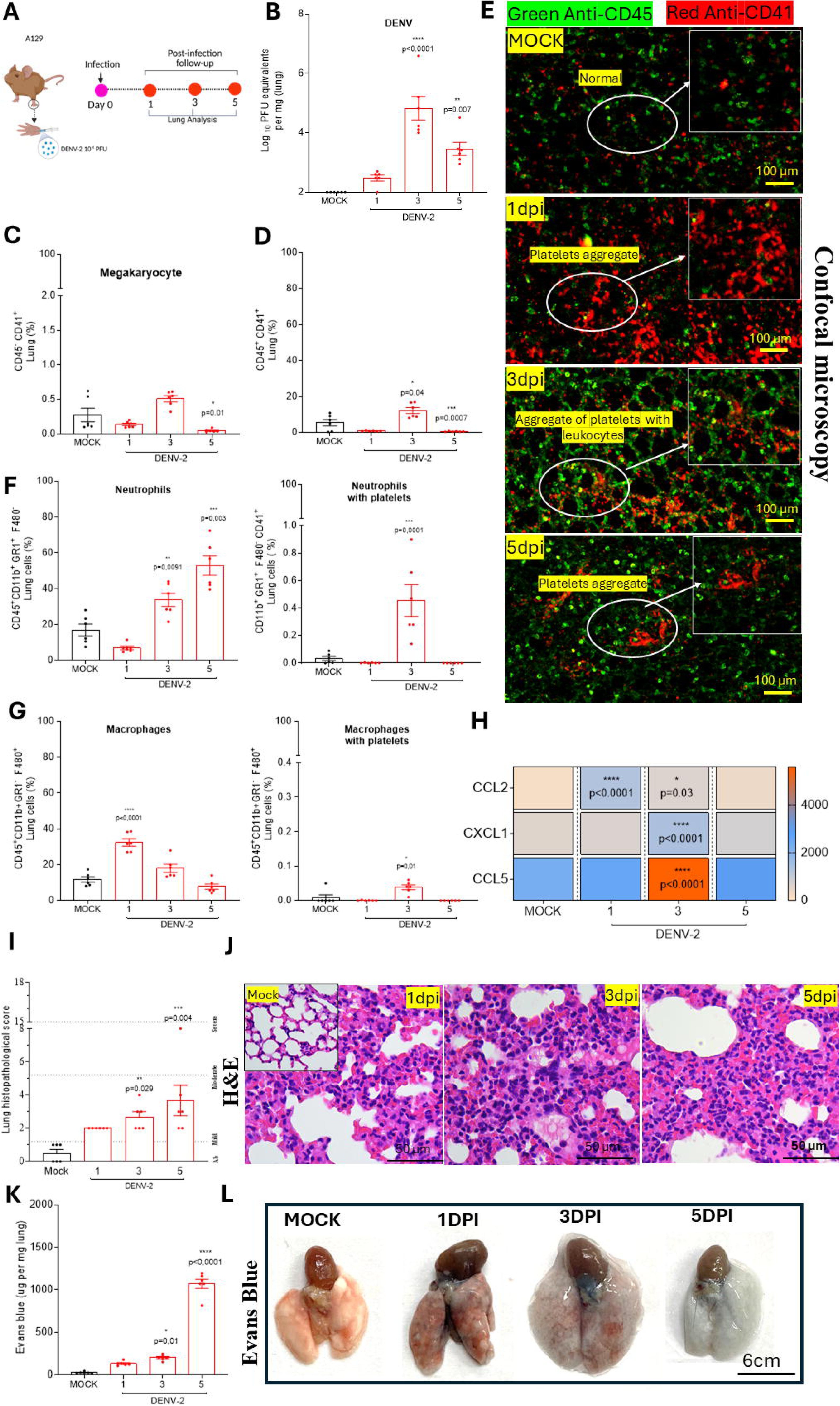
Assessment of Lung Damage Associated with Dengue Virus Infection. (A) Experimental design: A129 mice (n = 6 per group), were subcutaneously inoculated in the plantar region with DENV-2 at a concentration of 2×10LJ PFU/animal or with cell supernatant (Mock group). Mice were monitored daily for weight loss and clinical score and euthanized on days 1, 3, and 5 post-infections, followed by lung collection for analysis. **(B) Viral titer:** Determined from lung extracts of DENV-2-infected mice using a plaque assay. Results are expressed as log10 PFU per gram of tissue. **(C) Percentage of megakaryocytes:** Positively selected as CD45LJ CD41LJ cells using flow cytometry. **(D) Platelet-leukocyte aggregates:** Selected based on the CD45LJ CD41LJ cell population. **(E) Lung confocal microscopy:** Thirty minutes before euthanasia, animals received intravenous Anti-CD45 APC (green) and Anti-CD41 PE (red). **(F) Percentage of neutrophils**: Identified in the CD45LJ CD11bLJ GR1LJ F4/80LJ population, including aggregates with CD41LJ platelets. **(G) Percentage of macrophages:** Identified in the CD45LJ CD11bLJ GR1LJ F4/80LJ population, including aggregates with CD41LJ platelets. **(H) Heatmap of inflammatory mediators:** Quantification of CCL2, CXCL1, and CCL5 inflammatory mediators in the lung was performed by ELISA, with results expressed in pg/mg of lung tissue. **(I) Lung histopathological score. (J) Representative images of lung sections stained with H&E. (K) Assessment of vascular permeability in the lung:** Performed using the Evans blue dye technique. **(L) Representative images of the Evans blue technique in the lung**. The symbols (*, **, ***, ****) indicate statistically significant differences between infected groups and the control group (Mock) (p < 0.05). Statistical analysis: Figures 4E, I, J, and CCL2 levels: Kruskal-Wallis test followed by Dunn’s post-test. Figures 4B-F and L-P: One-way ANOVA followed by Dunnett’s post-test. Figure D: Statistical difference between the Mock and 5 dpi groups obtained using a t-test. Figure A was created with Biorender. Batista, VL (2025).

### 3.5 P2Y12 Receptor and P-Selectin Mediate Dengue Virus-Induced Thrombocytopenia

Our findings thus far demonstrate a significant increase in both soluble P-selectin levels and surface expression of P-selectin on platelets **(Figures 3C–E).** A Spearman correlation analysis revealed a negative Pearson correlation (r = –0.41, p =0.04) between platelet surface P-selectin expression in peripheral blood and total platelet count **(Suplementary Figure 4A)**, suggesting a potential role of P-selectin in dengue virus-induced thrombocytopenia. To further investigate this hypothesis, we assessed the role of P-selectin in DENV-induced thrombocytopenia. Upon platelet activation, P-selectin is rapidly translocated from α-granules to the cell surface, where it mediates the formation of platelet-leukocyte aggregate[16,33]. To evaluate the role of P-selectin in this context, we blocked P-Selectin function using a mouse-specific anti-CD62P monoclonal antibody **(Figure 5A).** Administration of the anti-P-selectin antibody to DENV-2–infected mice did not alter clinical scores (**Supplementary Figure 4B)**, nor did it affect the total number of megakaryocytes in the bone marrow **(Supplementary Figure 4C)**. In peripheral blood, P-selectin blockade prevented thrombocytopenia **(Figure 5B)**, reduced platelet activation and their aggregation with neutrophils and monocytes while preserving the total counts of these cells **(Figures 5C–F).**Treatment with anti-P-selectin also decreased platelet aggregates with dendritic cells **(Supplementary Figure 4D)** and led to decreased plasma levels of CXCL1 **(Figure 5G)** and MCPT-1 **(Suplementary Figure 4E).** In the lungs, P-selectin blockade reduced the total number of neutrophils and increased the number of macrophages **(Figure 5H-I)**. Moreover, it decreased platelet aggregates with neutrophils, macrophages, and dendritic cells **(Figure 5H-J)**, and lowered CXCL1 and CCL5 levels in the pulmonary environment **(Figure K).** These findings were associated with improved pulmonary histopathological scores **(Figures 5L-M).** Importantly, P-selectin inhibition did not affect viremia or viral load in the liver or lungs **(Supplementary 4F),** indicating that its blockade improves inflammatory and hematological parameters without impairing the ability of the host to deal with the infection. Our findings to date suggest that P-selectin contributes to dengue virus–induced thrombocytopenia.

**Figure 5:**
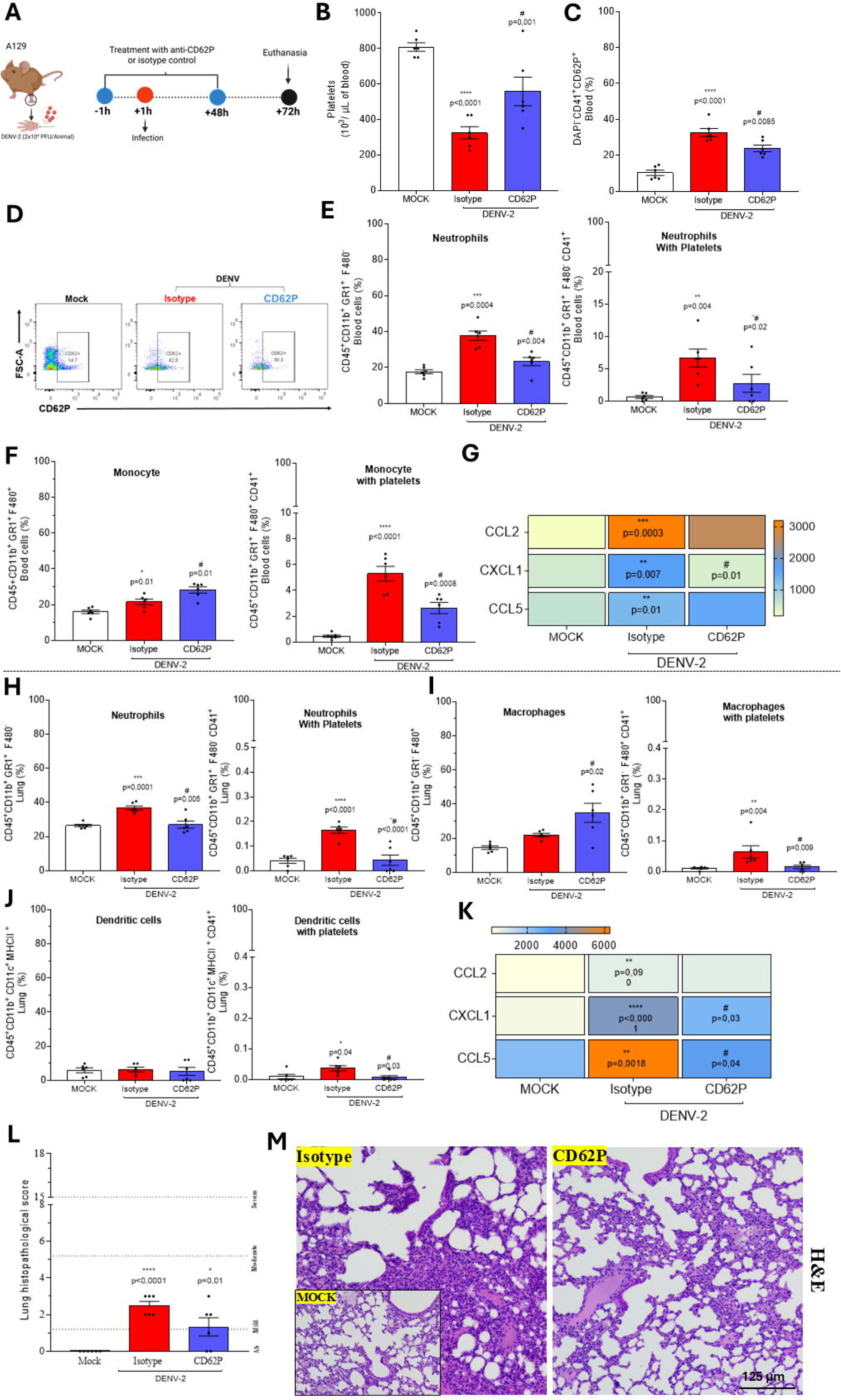
Evaluation of the Role of P-Selectin in Dengue Virus-Induced Thrombocytopenia. (A) Experimental Design. A129 mice (n = 6 per group) received an intraperitoneal injection of 200 µL containing 30 µg/animal of murine anti-CD62P antibody (CD62P group), murine anti-IgG antibody (Isotype), with cell supernatant (Mock group) one hour before infection. One hour later, mice were subcutaneously inoculated via the intraplantar route with 30 µL of DENV-2 at a concentration of 2×10LJ PFU/animal diluted in saline or with cell culture supernatant (Mock group). After 48 hours of infection, mice received additional injections with the same concentrations of anti-CD62P antibody, IgG, or saline. Mice were monitored daily for weight loss and clinical signs, and three days post-infection, were euthanized for organ collection for flow cytometry and inflammatory parameter analyses. **(B) Total Platelet Count in Peripheral Blood.** Performed manually using a Neubauer chamber, with results expressed as 10³ platelets/mL of blood. **(C) Percentage of CD62P-Positive Platelets in Peripheral Blood. (D) Representative dot plot of CD62P on the platelet surface.** For the assessment of platelet activation, platelets were identified as DAPILJ CD41LJ, and within this population, CD62P expression was evaluated. Platelets were analyzed on a logarithmic scale to exclude larger cells**. (E) Percentage of Neutrophils in Peripheral Blood.** Identified within the CD45+ CD11b+ GR-1+ F4/80+ population, including aggregates with CD41+ platelets. **(F) Percentage of Monocytes in Peripheral Blood.** Identified within the CD45+ CD11b+ GR-1+ F4/80+ population, including aggregates with CD41+ platelets. **(G) Plasma Inflammatory Mediator Heatmap.** CCL2, CXCL1, and CCL5 quantification was performed by ELISA, with results expressed in pg/mL. (H) **Percentage of Neutrophils in Lung.** Identified within the CD45+ CD11b+ GR-1+ F4/80+ population, including aggregates with CD41+ platelets. **(I) Percentage of Macrophages in the Lung.** Identified within the CD45+ CD11b+ GR-1^-^ F4/80+ population, including aggregates with CD41+ platelets. **(J) Percentage of Dendritic Cells in the Lung.** Identified within the CD45+ CD11b+ CD11c+ MHCII+ population, including aggregates with CD41+ platelets**. (K) Lung Inflammatory Mediator Heatmap.** CCL2, CXCL1, and CCL5 quantification was performed by ELISA, with results expressed in pg/mg of lung tissue. **(L) Lung Histopathological Score. (M) Representative H&E-Stained Lung Images.** Statistical significance is indicated as follows: Symbols (*, **, ***, ****) denote significant differences between infected groups treated with Isotype compared to the Mock group (p < 0.05). The symbol (#) indicates significant differences between the anti-CD62P antibody-treated group and the Isotype group (p < 0.05). Data that passed the normality test were analyzed using ANOVA followed by Dunnett’s post hoc test, comparing the groups to the isotype control. Data that did not pass the normality test were analyzed using the Kruskal-Wallis test followed by Dunn’s post hoc test. Figure A was created with Biorender. Batista, VL (2025).

Based on this, we next investigated the mechanism of platelet activation mediated by adenosine diphosphate (ADP) during dengue virus infection, a pathway that plays a critical role through its interaction with G protein–coupled receptors P2Y1 and P2Y12[34]. As a therapeutic strategy, we employed clopidogrel, a widely used antiplatelet agent in clinical practice **(Figure 6A).** Clopidogrel is a prodrug that, once metabolized in the liver, irreversibly inhibits the ADP receptor P2Y12, thereby suppressing platelet activation and reducing platelet-leukocyte interactions [24,35,36]. Our results indicate that treatment with clopidogrel prevented thrombocytopenia **(Figures 6B)** and reduced platelet activation **(Figures 6C-D)**, without affecting systemic platelet–leukocyte aggregation **(Figures 6E-F).** A decrease in plasma CCL5 levels was also observed **(Figure 6G).** In the lungs, clopidogrel treatment decreased platelet–leukocyte interaction, particularly with macrophages and dendritic cells **(Figures 6H–J).** Notably, clopidogrel did not alter clinical parameters in uninfected controls. These results show that platelet activation through ADP and the P2Y12 receptor lead to thrombocytopenia and thromboinflammation caused by dengue virus, with platelet surface P-selectin being a key factor in this mechanism. Additionally, administration of the antiplatelet agent clopidogrel demonstrated partial attenuation of infection-induced effects in a murine model of severe dengue.

**Figure 6:**
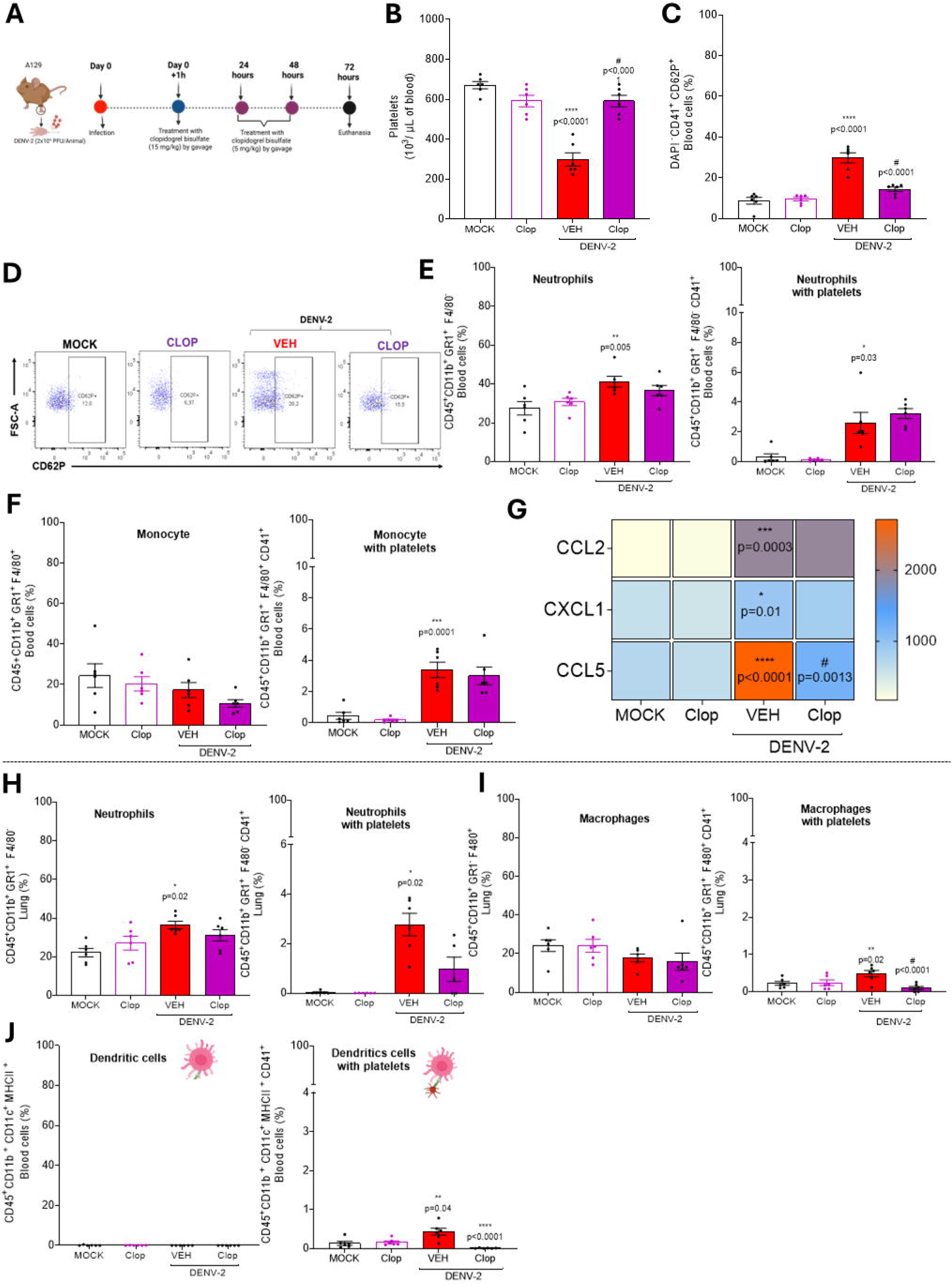
Assessment of the effects of clopidogrel administration on dengue virus infection. (A) **Experimental design**: A129 mice (n = 6 per group) were subcutaneously inoculated via the intraplantar route with 30 µL of DENV-2 (2×10LJ PFU/animal) diluted in saline or with cell supernatant (Mock group and Clop group). One hour after infection, mice were treated by gavage with an initial dose of 15 mg/kg of clopidogrel resuspended in 0.5% CMC, followed by a daily maintenance dose of 5 mg/kg under the same conditions. Mock and VEH groups received only the vehicle (0.5% CMC). The CLOP group (uninfected) received the same clopidogrel concentrations to assess drug toxicity**. (B) Total platelets count in peripheral blood**: Manually performed using a Neubauer chamber, with results expressed as 10³ platelets/mL of blood. (C) **Percentage of CD62P-positive platelets in peripheral blood**. (**D) Representative dot plot of the activated platelet population. (E) Percentage of Neutrophils in Peripheral Blood.** Identified within the CD45+ CD11b+ GR-1+ F4/80+ population, including aggregates with CD41+ platelets. **(F) Percentage of Monocytes in Peripheral Blood.** Identified within the CD45+ CD11b+ GR-1+ F4/80+ population, including aggregates with CD41+ platelets. **(G) Plasma Inflammatory Mediator Heatmap.** CCL2, CXCL1, and CCL5 quantification were performed by ELISA, with results expressed in pg/mL. (**H) Percentage of Neutrophils in Lung.** Identified within the CD45+ CD11b+ GR-1+ F4/80+ population, including aggregates with CD41+ platelets. **(I) Percentage of Macrophages in the Lung.** Identified within the CD45+ CD11b+ GR-1^-^ F4/80+ population, including aggregates with CD41+ platelets. **(J) Percentage of Dendritic Cells in the Lung.** Identified within the CD45+ CD11b+ CD11c+ MHCII+ population, including aggregates with CD41+ platelets. Symbols (*, **, ***, ****) denote significant differences between infected VEH and Mock groups (p < 0.05). Symbol # indicates significant differences between the clopidogrel-treated group and the infected VEH-treated group (p < 0.05). ANOVA with Dunnett’s post-test and Kruskal-Wallis with Dunn’s post-test were used, depending on the dataset.

## 4. DISCUSSION

Thrombocytopenia is a hallmark of dengue virus (DENV) infection and is strongly associated with disease severity and poor clinical outcomes [37]. In this study, using a model of severe dengue infection, we provide novel mechanistic insights into the pathogenesis of thrombocytopenia. Our findings demonstrate that DENV infection impairs thrombopoiesis and enhances platelet clearance, leading to thrombocytopenia and thromboinflammation through ADP–P2Y12 signaling and a key role for P-selectin.

We demonstrate that DENV targets the bone marrow, leading to a marked reduction in megakaryocyte numbers. These megakaryocytes are functionally impaired and undergo necrotic cell death. Injured megakaryocytes release platelets into the bone marrow microenvironment that exhibit features of apoptosis, including mitochondrial membrane depolarization and phosphatidylserine (PS) externalization. These alterations may contribute to platelet clearance and are consistent with the thrombocytopenia, platelet activation[38–41]and apoptotic features observed in the peripheral blood of dengue patients [42]. These findings align with previous *in vitro* studies showing that DENV impairs megakaryocyte polyploidization in megakaryocytic cell lines by modulating cell signaling pathways [43–45]. Another key component in the regulation of thrombopoiesis is thrombopoietin (TPO), a cytokine constitutively produced by the liver. Circulating TPO levels are modulated by the number of platelets and megakaryocytes through binding to the c-Mpl receptor. Studies have shown that following an acute drop in platelet production, TPO levels begin to rise within approximately 8 hours and peak around 24 hours [38–41] . In our model, we observed a significant increase in plasma TPO levels on the fifth day post-infection, coinciding with the lowest megakaryocyte count in the bone marrow and platelet count in the peripheral blood. This phenomenon is frequently reported in patients with thrombocytopenia and has been associated with severe forms of dengue, potentially serving as an early marker of disease severity [38,46].

In addition to impaired platelet production, DENV infection promotes platelet activation. In our model, platelet activation was evidenced by increased surface P-selectin expression and elevated soluble P-selectin levels in plasma. Plasma levels of NS1, which can activate platelets via TLR4 [47], increased in parallel with viremia and sP-selectin. Activated platelets formed aggregates with leukocytes in peripheral blood, in agreement with previous findings in dengue patients [15]. Platelet-leukocyte aggregates (PLAs) are formed via platelet P-selectin binding to PSGL-1 on leukocytes, followed by activation of integrins and other adhesion molecules [16,17]. In dengue, this interaction promotes monocyte activation and inflammasome assembly [48]. Furthermore, another study demonstrated that circulating platelet–monocyte complexes activate the AKTz/mTOR signaling pathway and glycolysis, leading to histone methylation and consequent changes in gene expression that limit monocyte responses to TLR agonists [49]. We observed increased plasma levels of CCL5, CXCL1, and CXCL4, chemokines secreted by activated platelets, further supporting the inflammatory role of platelets during DENV infection.

We also observed inflammatory responses in the lungs, with increased levels of inflammatory mediators and platelet-leukocyte aggregates in lung tissue. The lungs have been identified as sites of platelet production [31], and were recently identified as a major site of DENV infection in fatal pediatric cases [50]. Additionally, a retrospective study conducted at KS Hegde Hospital in Mangalore, India, analyzed 255 dengue cases and reported that approximately 10% of patients (n = 26) exhibited pulmonary manifestations, with pleural effusion being the most frequently observed complication [51]. In our model, pulmonary inflammation was more intense than hepatic inflammation, reinforcing that the lungs are target organs in dengue.

We hypothesize that P-selectin is a key mediator of the thromboinflammatory phenotype observed during dengue virus (DENV) infection. In our murine model, we demonstrate for the first time that P-selectin blockade significantly mitigates thrombocytopenia and reduce platelet activation. In addition, inhibition of P-selectin decreased platelet–leukocyte aggregates in both blood and lungs, underscoring its central role in DENV-induced thromboinflammation. Upregulated P-selectin expression has been associated with disease pathogenesis in several viral infections, including SARS-CoV-2 and Chikungunya [52–54]. In murine models of sickle cell disease, P-selectin promotes platelet–neutrophil aggregation and pulmonary inflammation [20,21]. Furthermore, both cellular and soluble forms of P-selectin, through interaction with PSGL-1, can induce NETosis, identifying this pathway as a potential therapeutic target in NET-mediated diseases [55]. In animal models of acute respiratory distress syndrome (ARDS), administration of monoclonal antibodies targeting P-selectin or Sialyl-Lewis-X, a natural PSGL-1 ligand, dramatically reduced lung injury [56]. In humans, elevated levels of soluble P-selectin have been detected in ARDS patients, particularly among non-survivors. Moreover, P-selectin antagonism has been shown to reduce macrophage accumulation in allografts during antibody-mediated rejection, further supporting its potential as a therapeutic target [56,57].

Given the thromboinflammatory phenotype observed in our model, we investigated the ADP-and P2Y12-mediated platelet activation pathway during dengue virus infection. For this purpose, we employed an antiplatelet therapeutic strategy using clopidogrel, a clinically used P2Y12 inhibitor. Clopidogrel treatment prevented thrombocytopenia and platelet activation, and reduced levels of inflammatory mediators in the plasma of infected mice. Additionally, treatment decreased platelet– leukocyte aggregate formation in the lungs.[24,35,58,59]. Beyond its antithrombotic effects, clopidogrel has demonstrated anti-inflammatory activity in cardiovascular and hypertensive models, where it reduces platelet–immune cell interactions and vascular inflammation. Similar thromboinflammatory mechanisms are shared across viral infections, such as SARS-CoV-2, in which clopidogrel-based combination therapy improved oxygenation and clinical outcomes in COVID-19 patients with prothrombotic profiles [24,35,58,59]. In the context of influenza-induced pneumonia, clopidogrel treatment also reduced platelet activation and platelet–leukocyte aggregate formation when combined with the antiviral oseltamivir. This combination therapy significantly improved animal survival, indicating a synergistic effect of clopidogrel with antiviral treatment.

In summary, this study provides the first *in vivo* demonstration, within a single model, of the impact of DENV infection on thrombopoiesis, platelet activation, and thromboinflammation. We identify the bone marrow and lungs as key target organs of the disease and show that P-selectin plays a central role in platelet–leukocyte interactions and dengue pathogenesis, suggesting that platelets are activated via ADP and the P2Y12 receptor during infection. These findings provide mechanistic insights and suggest potential therapeutic strategies for controlling dengue virus–induced thromboinflammation.

## Acknowledgements

The authors thank Ilma Marçal de Souza, Rosemeire Oliveira, and Tania Colina for their invaluable technical support. We also acknowledge the support of the Technological Center for Advanced and Innovative Therapies (CT-Terapias, UFMG) and the Experimental Platforms of UFMG, particularly the Flow Cytometry Facility at the Institutional Biomarker Research Laboratory (LINBIO) in the School of Pharmacy, UFMG.

## Authorship

**Contribution:** Batista VL, Costa VV, Queiroz-Junior CM, Martins MT, and Hottz ED conceived the study. Batista VL, Martins JR, Dias ASL, Silva LS, Fonseca TCM, Costa PAC, Silva WN, Pereira JAB, and Antunes M conducted the experiments. Guimarães PPG, Costa VV, Martins MT, and Menezes G provided financial support for the study. Batista VL interpreted the data, performed statistical analyses, and wrote the manuscript. All authors reviewed and approved the final version of the manuscript.

## Data sharing statement

For original data, please contact

---

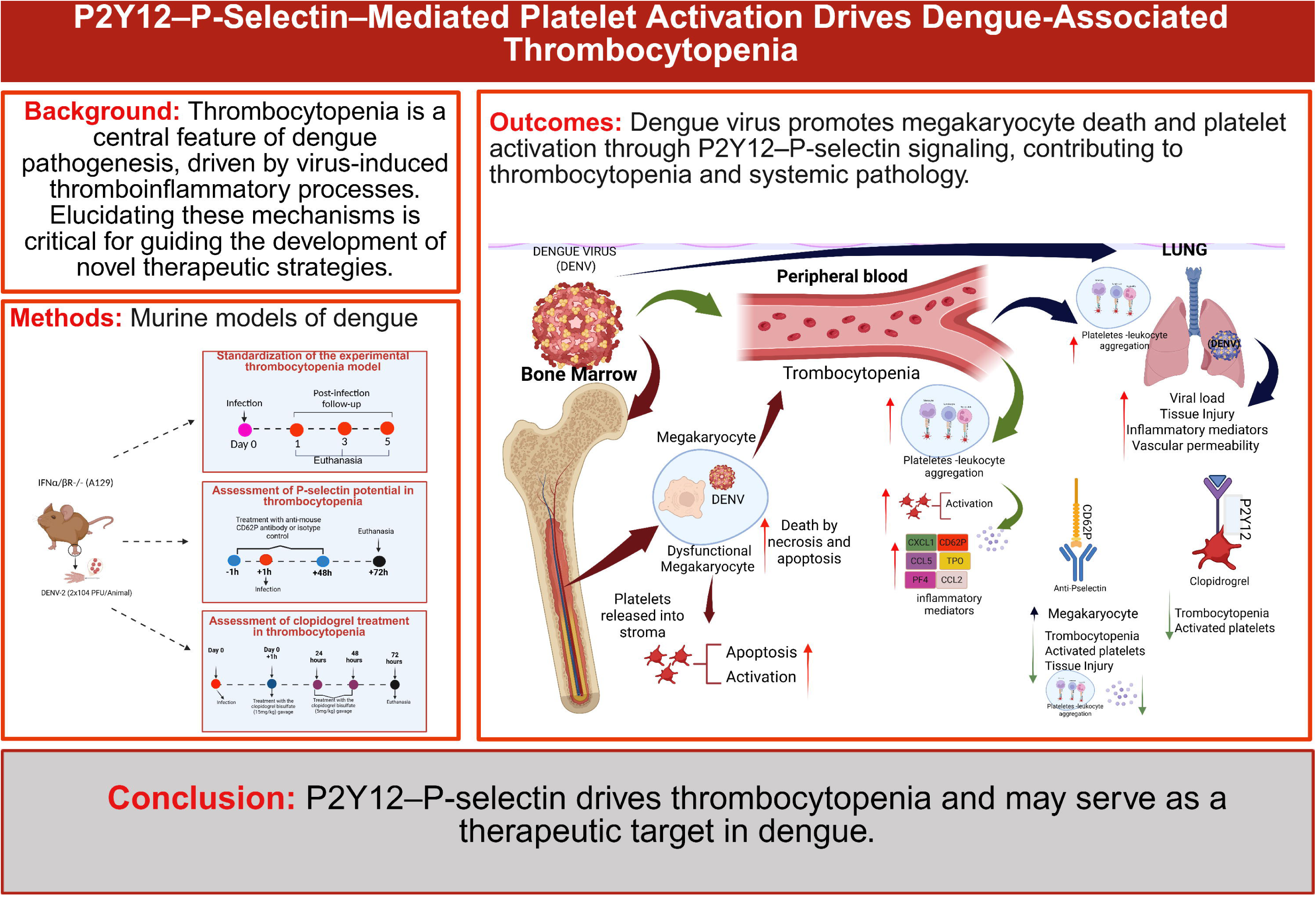

## REFERENCES

1. Guzman, M.G.; Gubler, D.J.; Izquierdo, A.; Martinez, E.; Halstead, S.B. Dengue Infection. Nat. Rev. Dis. Prim. 2016, 2, 1–26, doi:10.1038/nrdp.2016.55.

2. Haider, N.; Hasan, M.N.; Onyango, J.; Asaduzzaman, M. Global Landmark: 2023 Marks the Worst Year for Dengue Cases with Millions Infected and Thousands of Deaths Reported. IJID Reg. 2024, 13, 100459, doi:10.1016/j.ijregi.2024.100459.

3. Pan American Health Organization Report on the Epidemiological Situation of Dengue in the Americas. Paho/Who 2024, 582, 1–3.

4. Ojha, A.; Nandi, D.; Batra, H.; Singhal, R.; Annarapu, G.K.; Bhattacharyya, S.; Seth, T.; Dar, L.; Medigeshi, G.R.; Vrati, S.;, et al. Platelet Activation Determines the Severity of Thrombocytopenia in Dengue Infection. Sci. Rep. 2017, 7, 41697, doi:10.1038/srep41697.

5. Adane, T.; Getawa, S. Coagulation Abnormalities in Dengue Fever Infection: A Systematic Review and Meta-Analysis. PLoS Negl. Trop. Dis. 2021, 15, e0009666, doi:10.1371/journal.pntd.0009666.

6. Srichaikul, T.; Nimmannitya, S. Haematology in Dengue and Dengue Haemorrhagic Fever. Best Pract. Res. Clin. Haematol. 2000, 13, 261–276, doi:10.1053/beha.2000.0073.

7. BIERMAN, H.R. Hematodepressive Virus Diseases of Thailand. Ann. Intern. Med. 1965, 62, 867, doi:10.7326/0003-4819-62-5-867.

8. Vogt, M.B.; Lahon, A.; Arya, R.P.; Spencer Clinton, J.L.; Rico-Hesse, R. Dengue Viruses Infect Human Megakaryocytes, with Probable Clinical Consequences. PLoS Negl. Trop. Dis. 2019, 13, e0007837, doi:10.1371/journal.pntd.0007837.

9. Alonzo, M.T.G.; Lacuesta, T.L. V.; Dimaano, E.M.; Kurosu, T.; Suarez, L.C.; Mapua, C.A.; Akeda, Y.; Matias, R.R.; Kuter, D.J.; Nagata, S.;, et al. Platelet Apoptosis and Apoptotic Platelet Clearance by Macrophages in Secondary Dengue Virus Infections. J. Infect. Dis. 2012, 205, 1321–1329, doi:10.1093/infdis/jis180.

10. Michels, M.; Alisjahbana, B.; de Groot, P.G.; Indrati, A.R.; Fijnheer, R.; Puspita, M.; Dewi, I.M.W.; van de Wijer, L.; de Boer, E.M.S.; Roest, M.;, et al. Platelet Function Alterations in Dengue Are Associated with Plasma Leakage. Thromb. Haemost. 2014, 112, 352–362, doi:10.1160/TH14-01-0056.

11. Wills, B.A.; Oragui, E.E.; Stephens, A.C.; Daramola, O.A.; Dung, N.M.; Loan, H.T.; Chau, N.V.; Chambers, M.; Stepniewska, K.; Farrar, J.J.;, et al. Coagulation Abnormalities in Dengue Hemorrhagic Fever: Serial Investigations in 167 Vietnamese Children with Dengue Shock Syndrome. Clin. Infect. Dis. 2002, 35, 277–285, doi:10.1086/341410.

12. Alharbi, M.G.; Alanazi, N.; Yousef, A.; Alanazi, N.; Alotaibi, B.; Aljurf, M.; El Fakih, R. COVID-19 Associated with Immune Thrombocytopenia: A Systematic Review and Meta-Analysis. Expert Rev. Hematol. 2022, 15, 157–166, doi:10.1080/17474086.2022.2029699.

13. Hottz, E.D.; Lopes, J.F.; Freitas, C.; Valls-de-Souza, R.; Oliveira, M.F.; Bozza, M.T.; Da Poian, A.T.; Weyrich, A.S.; Zimmerman, G.A.; Bozza, F.A.;, et al. Platelets Mediate Increased Endothelium Permeability in Dengue through NLRP3-Inflammasome Activation. Blood 2013, 122, 3405–3414, doi:10.1182/blood-2013-05-504449.

14. Trugilho, M.R. de O.; Hottz, E.D.; Brunoro, G.V.F.; Teixeira-Ferreira, A.; Carvalho, P.C.; Salazar, G.A.; Zimmerman, G.A.; Bozza, F.A.; Bozza, P.T.; Perales, J. Platelet Proteome Reveals Novel Pathways of Platelet Activation and Platelet-Mediated Immunoregulation in Dengue. PLOS Pathog. 2017, 13, e1006385, doi:10.1371/journal.ppat.1006385.

15. Hottz, E.D.; Medeiros-de-Moraes, I.M.; Vieira-de-Abreu, A.; de Assis, E.F.; Vals-de-Souza, R.; Castro-Faria-Neto, H.C.; Weyrich, A.S.; Zimmerman, G.A.; Bozza, F.A.; Bozza, P.T. Platelet Activation and Apoptosis Modulate Monocyte Inflammatory Responses in Dengue. J. Immunol. 2014, 193, 1864–1872, doi:10.4049/jimmunol.1400091.

16. Furie, B.; Furie, B.C. Role of Platelet P-Selectin and Microparticle PSGL-1 in Thrombus Formation. Trends Mol. Med. 2004, 10, 171–178, doi:10.1016/j.molmed.2004.02.008.

17. Moore, K.L.; Stults, N.L.; Diaz, S.; Smith, D.F.; Cummings, R.D.; Varki, A.; McEver, R.P. Identification of a Specific Glycoprotein Ligand for P-Selectin (CD62) on Myeloid Cells. J. Cell Biol. 1992, 118, 445–456, doi:10.1083/jcb.118.2.445.

18. Peshkova, A.D.; Saliakhutdinova, S.M.; Sounbuli, K.; Selivanova, Y.A.; Andrianova, I.A.; Khabirova, A.I.; Litvinov, R.I.; Weisel, J.W. The Differential Formation and Composition of Leukocyte-Platelet Aggregates Induced by Various Cellular Stimulants. Thromb. Res. 2024, 241, 109092, doi:10.1016/j.thromres.2024.109092.

19. Gotardo, E.M.F.; Gushiken, L.F.S.; Brito, P.L.; Leonardo, F.C.; Torres, L.S.; Agoulnik, S.; Kovarik, J.; Bruederle, A.; Millholland, J.; Costa, F.F.;, et al. Neutralization of IL-1β and P-Selectin Inhibition Attenuate Organ Injury in Mice with Sickle Cell Disease. Blood 2023, 142, 2487–2487, doi:10.1182/blood-2023-181895.

20. Sullivan, V..; Hawley, A..; Farris, D..; Knipp, B..; Varga, A..; Wrobleski, S..; Thanapron, P.; Eagleton, M..; Myers, D..; Fowlkes, J..;, et al. Decrease in Fibrin Content of Venous Thrombi in Selectin-Deficient Mice. J. Surg. Res. 2003, 109, 1– 7, doi:10.1016/S0022-4804(02)00041-0.

21. Yokoyama, S.; Ikeda, H.; Haramaki, N.; Yasukawa, H.; Murohara, T.; Imaizumi, T. Platelet P-Selectin Plays an Important Role in Arterial Thrombogenesis by Forming Large Stable Platelet-Leukocyte Aggregates. J. Am. Coll. Cardiol. 2005, 45, 1280– 1286, doi:10.1016/j.jacc.2004.12.071.

22. Costa, V.V.; Sugimoto, M.A.; Hubner, J.; Bonilha, C.S.; Queiroz-Junior, C.M.; Gonçalves-Pereira, M.H.; Chen, J.; Gobbetti, T.; Libanio Rodrigues, G.O.; Bambirra, J.L.;, et al. Targeting the Annexin A1-FPR2/ALX Pathway for Host-Directed Therapy in Dengue Disease. Elife 2022, 11, doi:10.7554/eLife.73853.

23. Costa, V. V.; Fagundes, C.T.; Valadão, D.F.; Cisalpino, D.; Dias, A.C.F.; Silveira, K.D.; Kangussu, L.M.; Ávila, T. V.; Bonfim, M.R.Q.; Bonaventura, D.;, et al. A Model of DENV-3 Infection That Recapitulates Severe Disease and Highlights the Importance of IFN-γ in Host Resistance to Infection. PLoS Negl. Trop. Dis. 2012, 6, e1663, doi:10.1371/journal.pntd.0001663.

24. An, X.; Jiang, G.; Cheng, C.; Lv, Z.; Liu, Y.; Wang, F. Inhibition of Platelets by Clopidogrel Suppressed Ang II Induced Vascular Inflammation, Oxidative Stress, and Remodeling. J. Am. Heart Assoc. 2018, 7, doi:10.1161/JAHA.118.009600.

25. St. John, A.L. Influence of Mast Cells on Dengue Protective Immunity and Immune Pathology. PLoS Pathog. 2013, 9, e1003783, doi:10.1371/journal.ppat.1003783.

26. Costa, V. V.; Fagundes, C.T.; Valadão, D.F.; Ávila, T. V.; Cisalpino, D.; Rocha, R.F.; Ribeiro, L.S.; Ascenção, F.R.; Kangussu, L.M.; Junior, C.M.Q.;, et al. Subversion of Early Innate Antiviral Responses during Antibody-Dependent Enhancement of Dengue Virus Infection Induces Severe Disease in Immunocompetent Mice. Med. Microbiol. Immunol. 2014, 203, 231–250, doi:10.1007/s00430-014-0334-5.

27. Marques, P.E.; Antunes, M.M.; David, B.A.; Pereira, R. V; Teixeira, M.M.; Menezes, G.B. Imaging Liver Biology in Vivo Using Conventional Confocal Microscopy. Nat. Protoc. 2015, 10, 258–268, doi:10.1038/nprot.2015.006.

28. Pimenta, J.C.; Beltrami, V.A.; Oliveira, B. da S.; Queiroz-Junior, C.M.; Barsalini, J.; Teixeira, D.C.; de Souza-Costa, L.P.; Lima, A.L.D.; Machado, C.A.; Parreira, B.Z.S.G.;, et al. Neuropsychiatric Sequelae in an Experimental Model of Post-COVID Syndrome in Mice. Brain. Behav. Immun. 2025, 128, 16–36, doi:10.1016/j.bbi.2025.03.022.

29. Costa, V. V; Fagundes, C.T.; Valada, D.F.; Bonfim, M.R.Q.; Silveira, D.; Kangussu, L.M.; Thiago, V.A.; Dias, C.F.; Silva, A.; Bonaventura, D.; et al. A Model of DENV-3 Infection That Recapitulates Severe Disease and Highlights the Importance of IFN-c in Host Resistance to Infection. 2012, 6, 12–15, doi:10.1371/journal.pntd.0001663.

30. St. John, A.L.; Rathore, A.P.S.; Yap, H.; Ng, M.-L.; Metcalfe, D.D.; Vasudevan, S.G.; Abraham, S.N. Immune Surveillance by Mast Cells during Dengue Infection Promotes Natural Killer (NK) and NKT-Cell Recruitment and Viral Clearance. Proc. Natl. Acad. Sci. 2011, 108, 9190–9195, doi:10.1073/pnas.1105079108.

31. Lefrançais, E.; Ortiz-Muñoz, G.; Caudrillier, A.; Mallavia, B.; Liu, F.; Sayah, D.M.; Thornton, E.E.; Headley, M.B.; David, T.; Coughlin, S.R.;, et al. The Lung Is a Site of Platelet Biogenesis and a Reservoir for Haematopoietic Progenitors. Nature 2017, 544, 105–109, doi:10.1038/nature21706.

32. Livada, A.C.; McGrath, K.E.; Malloy, M.W.; Li, C.; Ture, S.K.; Kingsley, P.D.; Koniski, A.D.; Vit, L.A.; Nolan, K.E.; Mickelsen, D.;, et al. Long-Lived Lung Megakaryocytes Contribute to Platelet Recovery in Thrombocytopenia Models. J. Clin. Invest. 2024, 134, doi:10.1172/JCI181111.

33. Hottz, E.D.; Lopes, J.F.; Freitas, C.; Valls-de-Souza, R.; Oliveira, M.F.; Bozza, M.T.; Da Poian, A.T.; Weyrich, A.S.; Zimmerman, G.A.; Bozza, F.A.;, et al. Platelets Mediate Increased Endothelium Permeability in Dengue through NLRP3-Inflammasome Activation. Blood 2013, 122, 3405–3414, doi:10.1182/blood-2013-05-504449.

34. Offermanns, S. Activation of Platelet Function Through G Protein–Coupled Receptors. Circ. Res. 2006, 99, 1293–1304, doi:10.1161/01.RES.0000251742.71301.16.

35. Duarte, J.D.; Cavallari, L.H. Pharmacogenetics to Guide Cardiovascular Drug Therapy. Nat. Rev. Cardiol. 2021, 18, 649–665, doi:10.1038/s41569-021-00549-w.

36. Liu, Z.; Xiang, Q.; Mu, G.; Xie, Q.; Zhou, S.; Wang, Z.; Chen, S.; Hu, K.; Gong, Y.; Jiang, J.;, et al. Effectiveness and Safety of Platelet ADP–P2Y12 Receptor Inhibitors Influenced by Smoking Status: A Systematic Review and Meta Analysis. J. Am. Heart Assoc. 2019, 8, doi:10.1161/JAHA.118.010889.

37. Das, S.; Abreu, C.; Harris, M.; Shrader, J.; Sarvepalli, S. Severe Thrombocytopenia Associated with Dengue Fever: An Evidence-Based Approach to Management of Thrombocytopenia. Case Rep. Hematol. 2022, 2022, 1–3, doi:10.1155/2022/3358325.

38. Cardier, J.E.; Balogh, V.; Perez-Silva, C.; Romano, E.; Rivas, B.; Bosch, N.; Rothman, A.L. Relationship of Thrombopoietin and Interleukin-11 Levels to Thrombocytopenia Associated with Dengue Disease. Cytokine 2006, 34, 155–160, doi:10.1016/j.cyto.2006.04.002.

39. Emmons, R.V.B.; Reid, D.M.; Cohen, R.L.; Meng, G.; Young, N.S.; Dunbar, C.E.; Shulman, N.R. Human Thrombopoietin Levels Are High When Thrombocytopenia Is Due to Megakaryocyte Deficiency and Low When Due to Increased Platelet Destruction. Blood 1996, 87, 4068–4071, doi:10.1182/blood.v87.10.4068.bloodjournal87104068.

40. McCarty, J.M.; Sprugel, K.H.; Fox, N.E.; Sabath, D.E.; Kaushansky, K. Murine Thrombopoietin MRNA Levels Are Modulated by Platelet Count. Blood 1995, 86, 3668–3675, doi:10.1182/blood.v86.10.3668.bloodjournal86103668.

41. Stoffel, R.; Wiestner, A.; Skoda, R. Thrombopoietin in Thrombocytopenic Mice: Evidence against Regulation at the MRNA Level and for a Direct Regulatory Role of Platelets. Blood 1996, 87, 567–573, doi:10.1182/blood.V87.2.567.bloodjournal872567.

42. Hottz, E.D.; Oliveira, M.F.; Nunes, P.C.G.; Nogueira, R.M.R.; Valls-de-Souza, R.; Da Poian, A.T.; Weyrich, A.S.; Zimmerman, G.A.; Bozza, P.T.; Bozza, F.A. Dengue Induces Platelet Activation, Mitochondrial Dysfunction and Cell Death through Mechanisms That Involve DC-SIGN and Caspases. J. Thromb. Haemost. 2013, 11, 951–962, doi:10.1111/jth.12178.

43. Banerjee, A.; Tripathi, A.; Duggal, S.; Banerjee, A.; Vrati, S. Dengue Virus Infection Impedes Megakaryopoiesis in MEG-01 Cells Where the Virus Envelope Protein Interacts with the Transcription Factor TAL-1. Sci. Rep. 2020, 10, 19587, doi:10.1038/s41598-020-76350-5.

44. Basak, S.; Dutta, S.; Khanal, S.; Neelakanta, G.; Sultana, H. Dengue Virus Modulates Critical Cell Cycle Regulatory Proteins in Human Megakaryocyte Cells. Sci. Rep. 2025, 15, 19016, doi:10.1038/s41598-025-02640-5.

45. Kaur, J.; Rawat, Y.; Sood, V.; Periwal, N.; Rathore, D.K.; Kumar, S.; Kumar, N.; Bhattacharyya, S. Replication of Dengue Virus in K562-Megakaryocytes Induces Suppression in the Accumulation of Reactive Oxygen Species. Front. Microbiol. 2022, 12, doi:10.3389/fmicb.2021.784070.

46. Matondang, A.V.; Widodo, D.; Zulkarnain, I.; Rengganis, I.; Trihandini, I.; Inada, K.; Endo, S.; Suhendro The Correlation between Thrombopoietin and Platelet Count in Adult Dengue Viral Infection Patients. Acta Med. Indones. 2004, 36, 62–69.

47. Chao, C.-H.; Wu, W.-C.; Lai, Y.-C.; Tsai, P.-J.; Perng, G.-C.; Lin, Y.-S.; Yeh, T.-M. Dengue Virus Nonstructural Protein 1 Activates Platelets via Toll-like Receptor 4, Leading to Thrombocytopenia and Hemorrhage. PLOS Pathog. 2019, 15, e1007625, doi:10.1371/journal.ppat.1007625.

48. Barbosa-Lima, G.; Hottz, E.D.; de Assis, E.F.; Liechocki, S.; Souza, T.M.L.; Zimmerman, G.A.; Bozza, F.A.; Bozza, P.T. Dengue Virus-Activated Platelets Modulate Monocyte Immunometabolic Response through Lipid Droplet Biogenesis and Cytokine Signaling. J. Leukoc. Biol. 2020, 108, 1293–1306, doi:10.1002/JLB.4MA0620-658R.

49. Li, C.; Ture, S.K.; Nieves-Lopez, B.; Blick-Nitko, S.K.; Maurya, P.; Livada, A.C.; Stahl, T.J.; Kim, M.; Pietropaoli, A.P.; Morrell, C.N. Thrombocytopenia Independently Leads to Changes in Monocyte Immune Function. Circ. Res. 2024, 134, 970–986, doi:10.1161/CIRCRESAHA.123.323662.

50. Moragas, L.J.; Arruda, L.V.; Oliveira, L. de L.S.; Alves, F. de A.V.; Salomão, N.G.; da Silva, J.F.R.; Basílio-de-Oliveira, C.A.; Basílio-de-Oliveira, R.P.; Mohana-Borges, R.; Azevedo, C.G.;, et al. Detection of Viral Antigen and Inflammatory Mediators in Fatal Pediatric Dengue: A Study on Lung Immunopathogenesis. Front. Immunol. 2025, 16, doi:10.3389/fimmu.2025.1487284.

51. Keshav, L.B.K.A.; Malhotra, K.; Shetty, S. Lung Manifestation of Dengue Fever: A Retrospective Study. Cureus 2024, doi:10.7759/cureus.60655.

52. Gomes de Azevedo-Quintanilha, I.; Campos, M.M.; Teixeira Monteiro, A.P.; Dantas do Nascimento, A.; Calheiros, A.S.; Oliveira, D.M.; Dias, S.S.G.; Soares, V.C.; Santos, J. da C.; Tavares, I.; et al. Increased Platelet Activation and Platelet-Inflammasome Engagement during Chikungunya Infection. Front. Immunol. 2022, 13, doi:10.3389/fimmu.2022.958820.

53. Arce, N.A.; Cao, W.; Brown, A.K.; Legan, E.R.; Wilson, M.S.; Xu, E.-R.; Berndt, M.C.; Emsley, J.; Zhang, X.F.; Li, R. Activation of von Willebrand Factor via Mechanical Unfolding of Its Discontinuous Autoinhibitory Module. Nat. Commun. 2021, 12, 2360, doi:10.1038/s41467-021-22634-x.

54. Huang, C.; Wang, Y.; Li, X.; Ren, L.; Zhao, J.; Hu, Y.; Zhang, L.; Fan, G.; Xu, J.; Gu, X.;, et al. Clinical Features of Patients Infected with 2019 Novel Coronavirus in Wuhan, China. Lancet 2020, 395, 497–506, doi:10.1016/S0140-6736(20)30183-5.

55. Etulain, J.; Martinod, K.; Wong, S.L.; Cifuni, S.M.; Schattner, M.; Wagner, D.D. P-Selectin Promotes Neutrophil Extracellular Trap Formation in Mice. Blood 2015, 126, 242–246, doi:10.1182/blood-2015-01-624023.

56. Sakamaki, F.; Ishizaka, A.; Handa, M.; Fujishima, S.; Urano, T.; Sayama, K.; Nakamura, H.; Kanazawa, M.; Kawashiro, T.; Katayama, M. Soluble Form of P-Selectin in Plasma Is Elevated in Acute Lung Injury. Am. J. Respir. Crit. Care Med. 1995, 151, 1821–1826, doi:10.1164/ajrccm.151.6.7539327.

57. Valenzuela, N.M.; Hong, L.; Shen, X.-D.; Gao, F.; Young, S.H.; Rozengurt, E.; Kupiec-Weglinski, J.W.; Fishbein, M.C.; Reed, E.F. Blockade of P-Selectin Is Sufficient to Reduce MHC I Antibody-Elicited Monocyte Recruitment In Vitro and In Vivo. Am. J. Transplant. 2013, 13, 299–311, doi:10.1111/ajt.12016.

58. Viecca, M.; Radovanovic, D.; Forleo, G.B.; Santus, P. Enhanced Platelet Inhibition Treatment Improves Hypoxemia in Patients with Severe Covid-19 and Hypercoagulability. A Case Control, Proof of Concept Study. Pharmacol. Res. 2020, 158, 104950, doi:10.1016/j.phrs.2020.104950.

59. Kazui, M.; Nishiya, Y.; Ishizuka, T.; Hagihara, K.; Farid, N.A.; Okazaki, O.; Ikeda, T.; Kurihara, A. Identification of the Human Cytochrome P450 Enzymes Involved in the Two Oxidative Steps in the Bioactivation of Clopidogrel to Its Pharmacologically Active Metabolite. Drug Metab. Dispos. 2010, 38, 92–99, doi:10.1124/dmd.109.029132.

