## Supplementary material for "P2Y12–P-Selectin Mediated Platelet Activation Drives Dengue-Associated Thrombocytopenia": Suplemmetary Figure

**SUPPLEMENTARY FIGURES AND TABLES**

**Supplementary Figure 1**

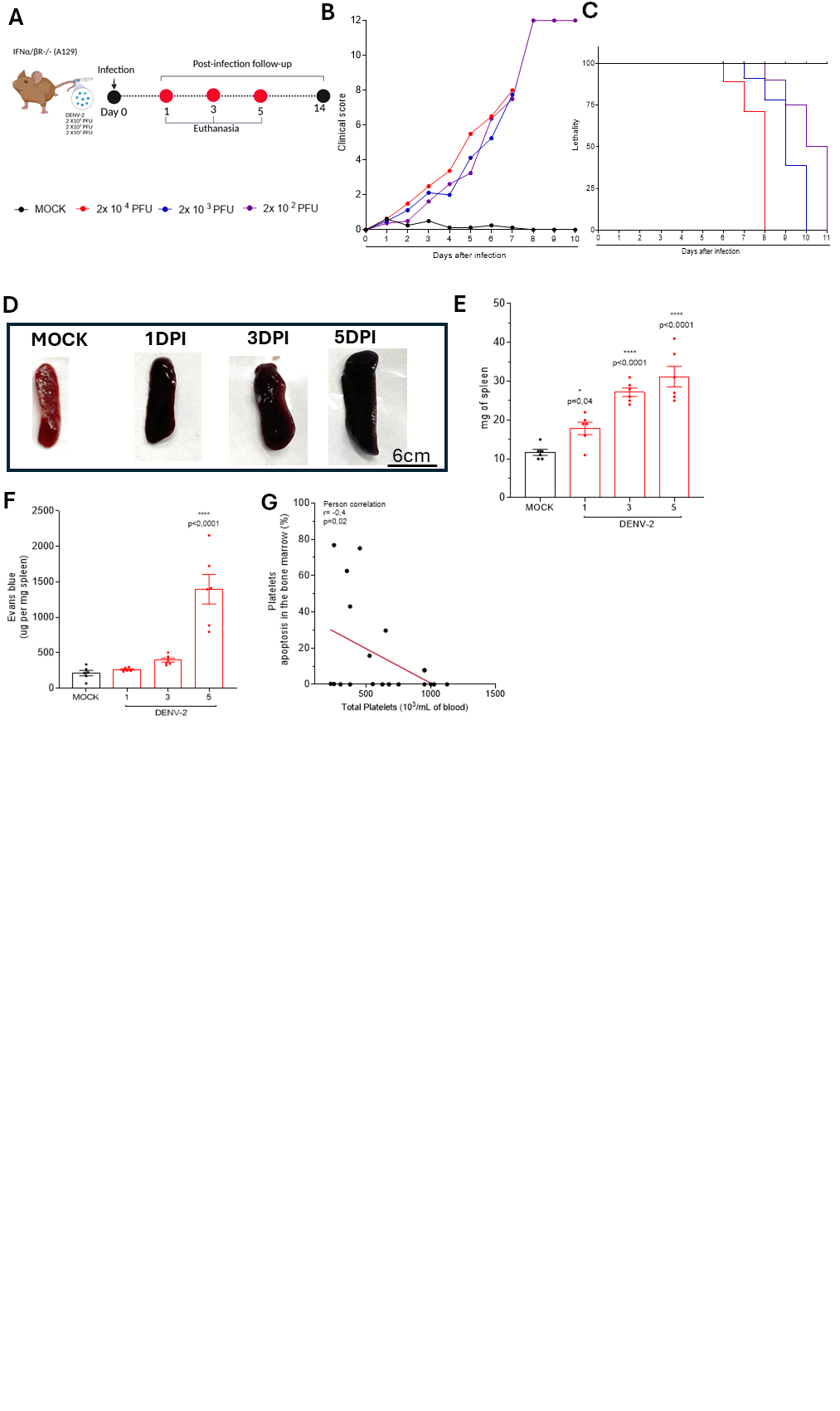

**Supplementary Figure 1: Standardization of the optimal inoculum to be administered to A129 mice via the intraplantar route. (A) Experimental scheme:** A129 mice were inoculated subcutaneously with three different concentrations of DENV-2 (2×10⁴ PFU, 2×10³ PFU, and 2×10³ PFU/animal) and monitored daily for 14 days for clinical score evaluation and weight loss**. (B) Kaplan-Meier survival curve (n = 4 per group). (C) Clinical score:** The animals were assessed daily for disease severity. The clinical score was determined based on the following parameters: weight loss, spinal hunching, diarrhea, ocular inflammation, decreased locomotor activity, and prostration. The score was assigned as detailed in **Table S2**. The graph shows the sum of the assigned scores. Animals with weight loss of 20% or more were euthanized (n = 4 per group**). (D) Spleen weight (mg). (E) Evaluation of vascular permeability in the spleen by the Evans blue technique. (F) Representative images of vascular permeability evaluation in the spleen.** In Figures D-F, animals were infected with 2×10⁴ PFU/animal or received 30 μL of saline solution and were euthanized on days 1, 3, and 5 post-infection. For vascular permeability evaluation, 100 μL of 0.5% Evans blue were injected intravenously 30 minutes before euthanasia. Animals were euthanized with an overdose of anesthetics, and a median laparotomy was performed with left ventricle perfusion with 1x PBS. Spleen, lung, and brain were collected for analysis. The amount of Evans blue remained in the tissue was determined by comparing the absorbance of the sample with a standard curve of Evans blue, read at 620 nm on a plate reader spectrophotometer. Results were expressed as the amount of Evans blue per 100 mg of tissue (n = 6). Symbols (*, **, ***, ****) indicate statistically significant differences between infected groups and the control group (Mock) (p < 0.05), as determined by one-way ANOVA followed by Dunnett's post-test.

**Supplementary Figure 2**

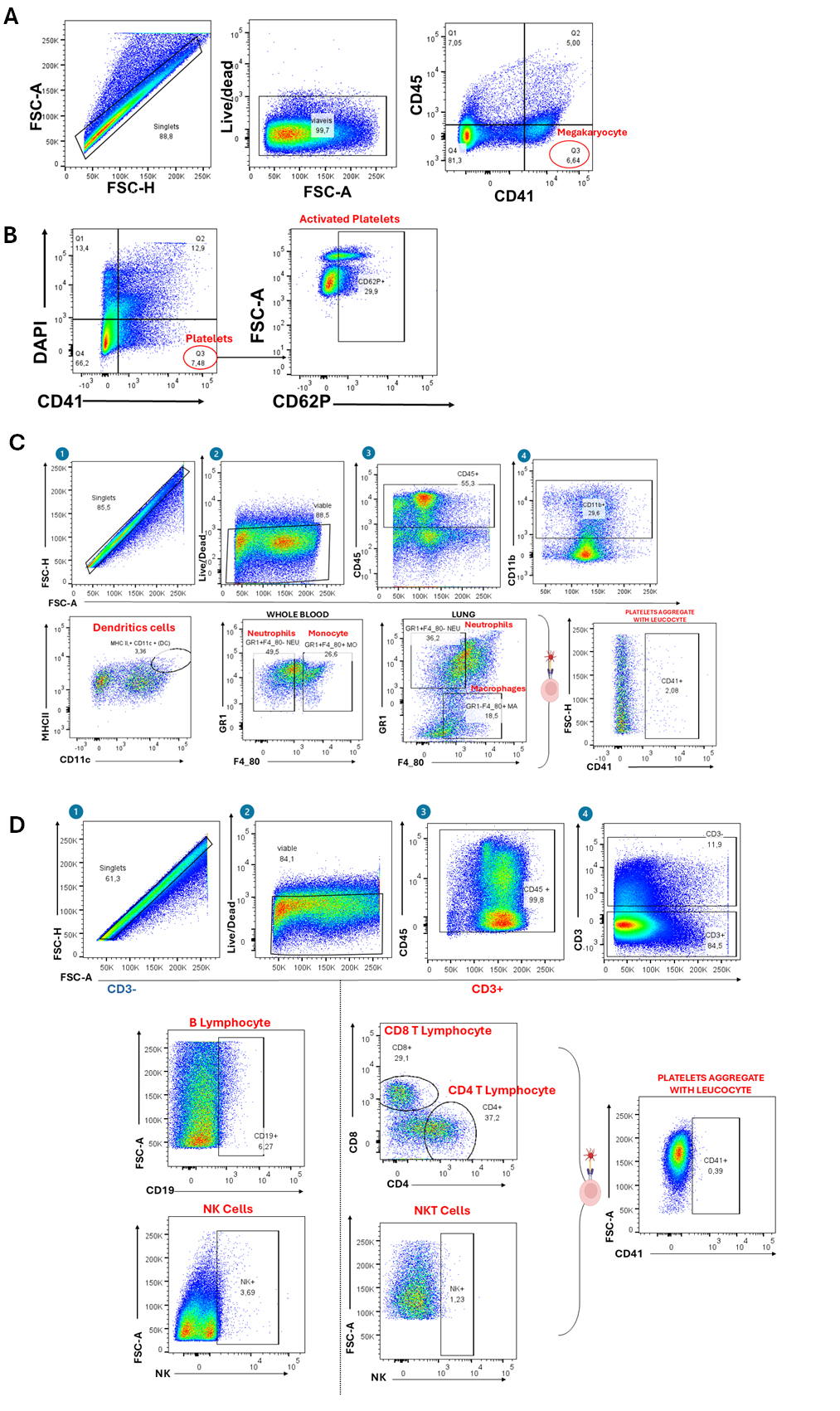

**Supplementary Figure 2: Gating strategy of the cell subsets evaluated in this study.
(A)** **Illustrative dot plots of the megakaryocyte subpopulation in the bone marrow:** first, single cells were selected (FSC-A × FSC-H), followed by the viable cell population. Within this population, megakaryocytes were identified as CD45⁻ and CD41⁺ cells. **(B) Representative density plots which illustrate the gating strategy for total platelets and activated platelets.** The same strategy was applied to bone marrow, blood, and lung samples. The cytometer readings were conducted using a logarithmic scale, excluding cellular dendrites and larger cells such as megakaryocytes and leukocytes. Platelets were identified as those labeled with DAPI-, due to the absence of nuclei, and CD41+. Platelet activation was characterized by the expression of CD62P.***.**  **(B) Representative density plots illustrating the gating strategy for the myeloid panel. First, gating for singlet cells (FSC-H x FSC-A) was performed.** Within this population, only viable cells were selected using LIVE/DEAD staining. From the viable cells, the population was strongly positive for CD45 was selected. Within this population, cells positive for CD11b were further sorted, and dendritic cells were identified as those positive for CD45, CD11b, CD11c, and MHCII. Macrophages were selected as CD45+, CD11b+ GR1- F4/80+, monocytes as CD45+ CD11b+ GR1+ F4/80+, and neutrophils as CD11b+ GR1+ F4/80-. Within each subset, additional gating was performed to identify CD41+ cells. **(C) Representative density plots illustrating the gating strategy for the lymphoid panel.** Initially, gating for singlet cells (FSC-H x FSC-A) was performed. Within this population, only viable cells, marked with LIVE/DEAD dye, were selected. From the viable cells, the population strongly positive for CD45 was chosen. CD3-positive cells were subdivided into CD4+ T cells, CD8+ T cells, and NKT cells. CD3-negative cells were subdivided into B lymphocytes and NK cells. For each cell subtype, CD41 staining was performed to identify platelet-leukocyte aggregates.

**Supplementary Figure 3**

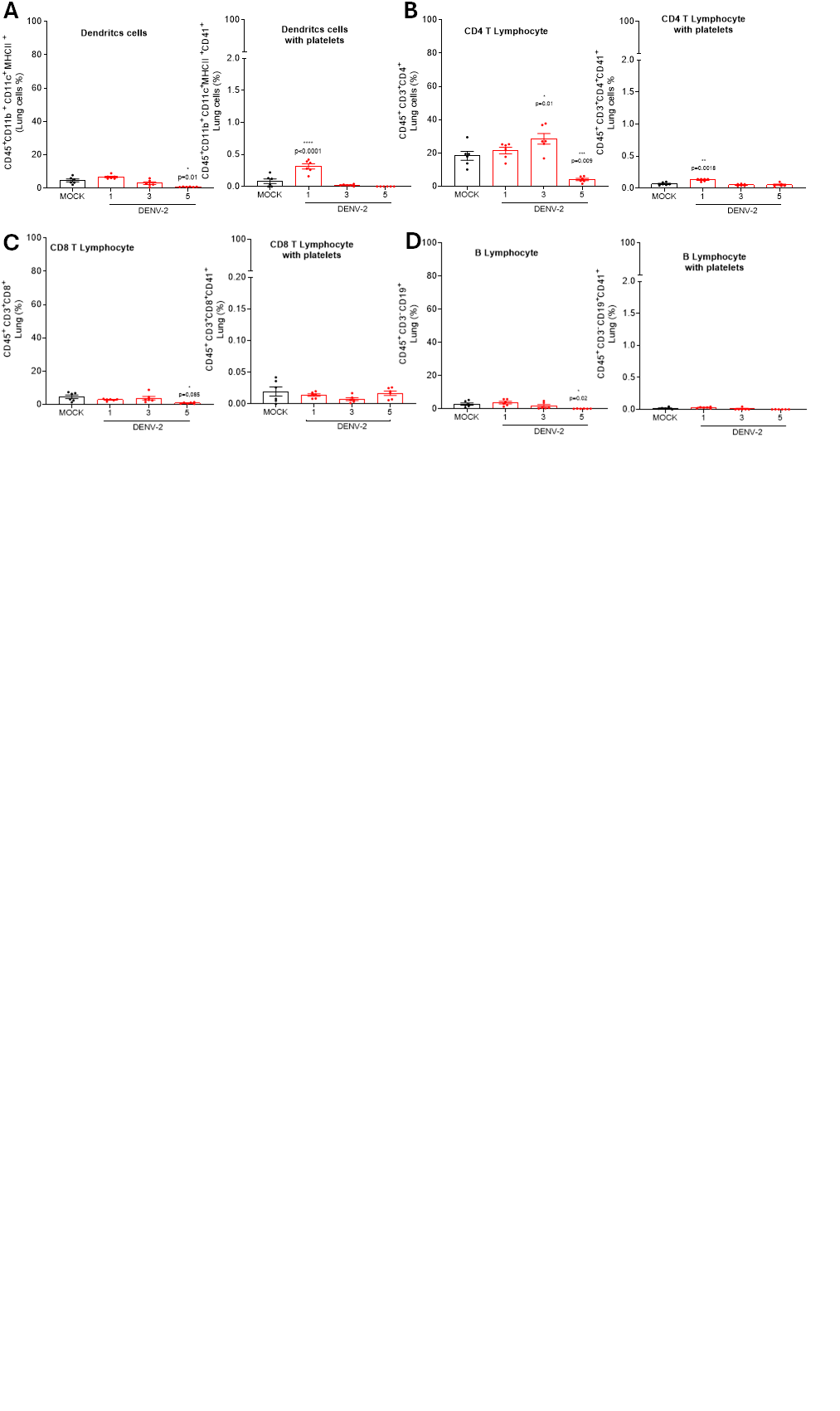

**Supplementary Figure 3:** **Complementary results of the other cellular subtypes from the pulmonary macerate**. (**A) Dendritic cells:** Selected from the CD45⁺ CD11b^+^ CD11c^+^ MHC II^+^ population and aggregated with CD41⁺ platelets. **(B) CD4⁺ T lymphocytes:** Selected from the CD45⁺ CD3⁺ CD4⁺ population and aggregated with CD41⁺ platelets**. (C) CD8⁺ T lymphocyte**s: Selected from the CD45⁺ CD3⁺ CD8⁺ population and aggregated with CD41⁺ platelets. **(D) B lymphocytes**: Selected from the CD45⁺ CD3⁻ CD19⁺ population and aggregated with CD41⁺ platelets. Symbols (*, **, ***, ****) indicate statistically significant differences between infected groups and the control group (Mock) (p < 0.05), as determined by one-way ANOVA followed by Dunnett’s post-test.

**Supplementary Figure 4**

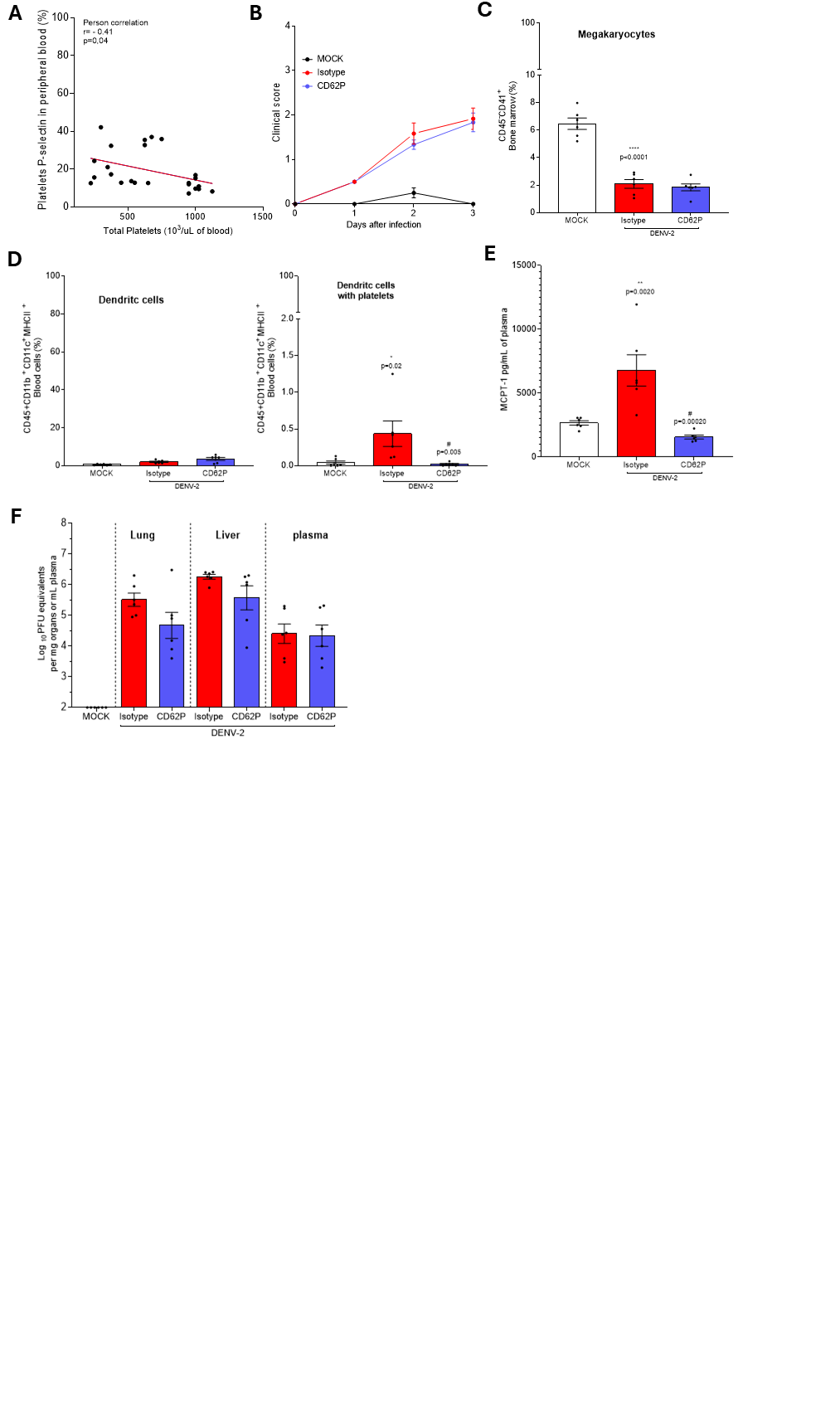

**Supplementary Figure 4: Supplementary data from the experiment evaluating P-selectin blockade in dengue virus infection. (A) Pearson correlation.** between manually counted platelets in peripheral blood and the percentage of CD62P expressed on the platelet surface**. (B) Clinical score:** The animals were assessed daily for disease severity. The clinical score was determined based on the following parameters: weight loss, spinal hunching, diarrhea, ocular inflammation, decreased locomotor activity, and prostration. The score was assigned based on severity: 0 (absence of symptoms), 0.5 (mild), 1 (moderate), and 2 (severe). The graph shows the sum of the scores assigned, as explained in Table S2. (**C) Percentage of megakaryocytes in the bone marrow.** Identified as the CD45⁻ CD41⁺ population. **(D) Percentage of dendritic cells in peripheral blood**. Identified as the CD45⁺ CD11b⁺ CD11c⁺ MHCII⁺ population, positive for CD41⁺ and aggregated with platelets. **(E) Plasma levels of MCPT-1**. were measured using the ELISA technique (**F) Viral Titer.** Determined in organ extracts (paw, spleen, liver, and brain) and plasma of DENV-2-infected mice using a plaque assay. Results are expressed as Log₁₀ PFU per gram of tissue or per milliliter of plasma

**TABLES**

**Table S1. Antibodies list for flow cytometry.**

| Antigen | [Fluorochrome](http://www.fluorochrome.com/) | Clone | Concentration | Company |
| --- | --- | --- | --- | --- |
| CD8a | eFluor 450 | 53-6.7 | 1/1300 | ThermoFisher |
| LIVE/DEAD | Acqua |  | 1/1000 | ThermoFisher |
| Streptavidin | Pacific Orange |  | 1/200 | ThermoFisher |
| CD45 | PercpCy5.5 | Ly-5.2 | 1/200 | BioLegend |
| CD11b | Super Bright 600 | M1/70 | 1/500 | ThermoFisher |
| CD11c | Super Bright 645 | N418 | 1/500 | ThermoFisher |
| CD45 | PerCP-Cy5.5 | Ly-5.2 | 1/200 | BioLegend |
| NK | PerCP-Cy5.5 | PK136 | 1/150 | ThermoFisher |
| CD4 | PE-Cyanine7 | GK1.5 | 1/3000 | ThermoFisher |
| F4/80 | APC | BM8 | 1/200 | ThermoFisher |
| MHC-II | APC-eFluor 780 | M5/114.15.2 | 1/500 | ThermoFisher |
| Gr-1 | Biotin | RB6-8C5 | 1/500 | BioLegend |
| CD45 | Pacific Orange | 30-F11 | 1/200 | ThermoFisher |
| CD41 | FITC | MLU Reg30 | 1/200 | BioLegend |
| CD41 | PE |  | 1/200 | Invitrogen |
| CD62P | PerCP-eFluor™ 710 | Psel.KO2.3 | 1/100 | eBioscience |

**Table S2. Clinical Scoring Table for Mice**

| Parameter | Score 0 | Score 0.5 | Score 1 | Score 1.5 | Score 2 |
| --- | --- | --- | --- | --- | --- |
| Posture | Normal | Slightly arched when at rest | Clearly hunched posture | Severely hunched, limited mobility | Severely hunched or immobile |
| Weight loss (% initial) | None | 1-5% | >5-10% | 11-15% | >15% |
| Conjunctivitis | Absent | Mild redness | Watery eyes | Partially closed eyes, moderate inflammation | Closed eyes or purulent discharge |
| Diarrhea | Normal | Slightly soft stools | Loose stools/diarrhea | Diarrhea with soiling of the perianal area | Severe diarrhea, extensive soiling |
| Locomotor activity | Normal | Slight reduction in movement | Noticeably reduced movement | Moves only when stimulated | Nearly or completely immobile |
| Pelage | Normal | little goosebumps | Slightly goosebumps | visibly goosebumps | dirty, frizzy hair |
| Animals are evaluated daily for the appearance of any symptoms listed in the table. The minimum score is 0 and the maximum is 12 points. Animals that exhibit a weight loss equal to or greater than 20% are euthanized, following the guidelines of humane endpoint criteria. Animals infected with the dengue virus are assessed in comparison with healthy control animals that did not receive the virus. | | | | | |
